# The *Legionella*-driven PtdIns(4)*P* gradient at LCV-ER membrane contact sites promotes Vap-, OSBP- and Sac1-dependent pathogen vacuole remodeling

**DOI:** 10.1101/2022.06.17.496549

**Authors:** Simone Vormittag, Dario Hüsler, Ina Haneburger, Tobias Kroniger, Aby Anand, Manuel Prantl, Caroline Barisch, Sandra Maaß, Dörte Becher, François Letourneur, Hubert Hilbi

## Abstract

The causative agent of Legionnaires’ disease, *Legionella pneumophila*, governs interactions with host cells by secreting ca. 330 different “effector” proteins. The facultative intracellular bacteria replicate in macrophages and amoeba within a unique compartment, the *Legionella*-containing vacuole (LCV). Hallmarks of LCV formation are the phosphoinositide (PI) lipid conversion from PtdIns(3)*P* to PtdIns(4)*P*, fusion with endoplasmic reticulum (ER)-derived vesicles and a tight association with the ER. Proteomics of purified LCVs revealed the presence of membrane contact sites (MCS) proteins implicated in lipid exchange. Using dually fluorescence-labeled *Dictyostelium discoideum* amoeba, we reveal that the VAMP-associated protein (Vap), the PtdIns(4)*P* 4-phosphatase Sac1, and the large fusion GTPase Sey1/atlastin-3 localize to the ER, but not to the LCV membrane, and that these ER-resident proteins promote intracellular replication of *L. pneumophila* and LCV remodeling. Moreover, oxysterol binding proteins (OSBPs) preferentially localize to the ER (OSBP8) or the LCV membrane (OSBP11), respectively, and promote (OSBP8) or restrict (OSBP11) intracellular replication of *L. pneumophila* and LCV expansion. Furthermore, the PtdIns(4)*P*-subverting *L. pneumophila* effectors LepB and SidC also promote LCV remodeling. Taken together, the *Legionella*- and host cell-driven PtdIns(4)*P* gradient at LCV-ER MCSs promotes Vap-, OSBP- and Sac1-dependent pathogen vacuole remodeling.

## Introduction

*Legionella pneumophila* is an amoeba-resistant environmental bacterium, which causes a severe pneumonia called Legionnaires’ disease [1, 2]. The facultative intracellular pathogen employs a conserved mechanism to grow in free-living protozoa as well as in alveolar macrophages, which is a prerequisite to cause disease [3–6]. To govern the interaction with host cells, *L. pneumophila* employs the Icm/Dot (Intracellular multiplication/defective organelle trafficking) type IV secretion system (T4SS), which translocates ca. 330 different “effector” proteins into host cells [6, 7]. These effector proteins subvert pivotal host cell processes, including the endocytic, secretory, retrograde and autophagy pathways, lipid metabolism, transcription, translation, and apoptosis [8–12].

Within host cells, *L. pneumophila* forms a unique, membrane-bound replication compartment, the *Legionella*-containing vacuole (LCV), which restricts interactions with the endocytic pathway [13–16]. Hallmarks of LCV formation are the phosphoinositide (PI) lipid conversion from phosphatidylinositol 3-phosphate (PtdIns(3)*P*) to PtdIns(4)*P*, the subversion of the small GTPase Rab1, fusion of the pathogen vacuole with endoplasmic reticulum (ER)-derived vesicles, and finally, a tight association with the ER [13–16]. The LCV is decorated with PtdIns(4)*P* [17], and PI conversion from the endocytic PI PtdIns(3)*P* to the secretory PI PtdIns(4)*P* occurs in the first 1-2 hours after bacterial uptake [18].

The production of PtdIns(4)*P* on the LCV membrane implicates the sequential activity of *L. pneumophila* Icm/Dot-secreted effector proteins: PtdIns 3-kinase MavQ [19], PtdIns(3)*P* 4-kinase LepB [20] and PtdIns(3,4)*P*_2_ 3-phosphatase SidF [21]. Moreover, host PI-modulating enzymes such as the PtdIns 4-kinases PI4KIIIα [22] and PI4KIIIβ [23], and the PtdIns(4,5)*P* 5-phosphatase OCRL1 [24] might also play a role in the production of PtdIns(4)*P* on the LCV membrane.

The small GTPase Rab1 is enriched in the *cis*-Golgi apparatus and controls ER-Golgi secretory trafficking [14]. Rab1 is recruited to the LCV membrane and activated by the Icm/Dot secreted effector SidM (*alias* DrrA), which binds to PtdIns(4)*P* through the novel P4M domain [22, 23, 25–27] and possess Rab1 guanine nucleotide exchange factor (GEF)/ guanine dissociation inhibitor (GDI) displacement factor (GDF) activity [28–34] as well as adenosine monophosphorylation (AMPylation) activity [35, 36]. Moreover, Rab1 is de-AMPylated by the effector SidD [37, 38] and reversibly phosphocholinated by the effectors AnkX and Lem3 [39–41]. The covalent modifications of GTP-bound Rab1 lock the small GTPase in its active state. Finally, Rab1 is inactivated by the GTPase activating protein (GAP) LepB [31]. The small GTPase ARF1 controls also retrograde Golgi-ER trafficking [14]. ARF1 is recruited to and activated on the LCV by the effector RalF, the first *L. pneumophila* GEF identified [42].

The LCV intercepts trafficking at ER exit sites between the ER and the Golgi apparatus, and inhibition of ER-Golgi transport blocks the formation of the pathogen vacuole [42–44]. Sec22b, a SNARE (soluble *N*-ethylmaleimide-sensitive factor (NSF) attachment protein (SNAP) receptor), which promotes fusion of ER-derived vesicles with the *cis*-Golgi, is recruited to LCVs shortly after infection [45, 46]. The activation of Rab1 by SidM on the LCV mediates the fusion of ER-derived vesicles with the pathogen vacuole through the non-canonical pairing of the vesicle (v-)SNARE Sec22b on the vesicles with plasma membrane (PM)-derived target (t-)SNAREs such as syntaxin (Stx) 2, 3, 4 and SNAP23 on the LCV [47, 48]. The LCV intercepts anterograde as well as retrograde trafficking between the ER and the Golgi apparatus [9, 11, 13, 14]. Accordingly, nascent LCVs continuously capture and accumulate PtdIns(4)*P*-positive vesicles from the Golgi apparatus, and their sustained association requires a functional T4SS [49]. LCVs also sequentially recruit the small GTPases Rab33b, Rab6a and the ER-resident SNARE syntaxin18 (Stx18), which are implicated in Golgi-ER retrograde trafficking, to promote the association of the pathogen vacuole with the ER [50].

The association of the LCV with ER in macrophages and amoeba has been initially documented more than 25 years ago [51, 52]. The vacuole containing *L. pneumophila* associates with rough ER within minutes after uptake, and dependent on the Icm/Dot T4SS, the vacuole and the ER form extended contact sites (> 0.5 µm in length), which are connected by tiny, periodic “hair-like” structures [53]. Remarkably, the ER elements remain attached to LCVs even after cell homogenization [53], and intact LCVs purified from *L. pneumophila*-infected *D. discoideum* co-purify with extensive fragments of calnexin-GFP-positive ER [54–56]. In dually fluorescence labeled *D. discoideum*, LCVs accumulate calnexin-positive ER within minutes, and the ER remains separate from the PtdIns(4)*P*-positive pathogen vacuole for at least 8 h post infection [18, 57]. The ER tubules interacting with LCVs are decorated with reticulon 4 (Rtn4) and the large fusion GTPase atlastin/Sey1, which are implicated in the architecture and dynamics of the ER, respectively [58, 59]. The *L. pneumophila* effector Ceg9 might stabilize the LCV-ER contact sites by directly binding to the host protein Rtn4 [58]. Finally, depletion of the PtdIns(4)*P* 4-phosphatase Sac1, a crucial enzyme of ER membrane contact sites (MCS), reduced the recruitment of endogenous Rab1 to LCVs [22].

The MCS between two distinct cellular compartments or organelles adopt a number of functions and are of crucial importance for the non-vesicular exchange of lipids in mammalian cells [60]. MCS between the ER and the PM or organelles promote lipid exchange driven by a gradient of PtdIns(4)*P*, which is established by PtdIns 4-kinases on the PM or organelles and maintained by the integral ER membrane protein PtdIns(4)*P* 4-phosphatase Sac1 [61, 62]. Driven by the PtdIns(4)*P* gradient, phosphatidylserine accumulates at the inner leaflet of the PM [63, 64] and cholesterol accumulates at the Golgi apparatus [61, 65]. The lipid exchange is promoted by lipid transfer proteins termed oxysterol binding proteins (OSBPs), several of which bind PtdIns(4)*P* through a PH domain and by the ER-resident Vap (vesicle-associated membrane protein (VAMP)-associated protein) through a FFAT (two phenylalanines (FF) in an acidic tract) motif [60].

Given the accumulation of PtdIns(4)*P* on the limiting LCV membrane and the tight association of the pathogen vacuole with ER, the LCV-ER interface might actually form functionally important MCS, which promote pathogen vacuole remodeling through a PtdIns(4)*P* gradient. Using dually fluorescence-labeled *D. discoideum* amoeba, we identify Vap, Sac1 and Sey1 on the ER, but not on the LCV membrane, while OSBP8 and OSBP11 exclusively localize to the ER or the LCV membrane, respectively. The MCS components are implicated in LCV remodeling and intracellular growth of *L. pneumophila*, and a LCV-to-ER PtdIns(4)*P* gradient is established by bacterial effector proteins. These findings indicate that the *Legionella*-driven PtdIns(4)*P* gradient at LCV-ER MCSs promotes Vap-, Sac1- and OSBP-dependent pathogen vacuole remodeling.

## Results

### Comparative proteomics of LCVs from *D. discoideum* wild-type and Δ*sey1* reveals MCS components

Given the tight and stable association of the LCV limiting membrane with the ER, MCS are likely formed between the compartments. The ER tubule-resident large GTPase Sey1/Atl3 (DDB_G0279823) promotes the decoration of LCVs with ER, and in absence of the GTPase, significantly less ER accumulates on the pathogen vacuole, while the PtdIns(4)*P*-positive LCV appears to form normally [59, 66]. To gain further insights into the process of LCV formation, and in particular the role of the ER, we analyzed by comparative proteomics LCVs from the *L. pneumophila*-infected *D. discoideum* Ax3 wild-type strain or the Δ*sey1* mutant. This approach revealed 3658 host or bacterial proteins identified on LCVs from *D. discoideum* Ax3 or Δ*sey1* (**Table S1**).

These proteins include the OSBPs OSBP7 (*osbG*) and OSBP8 (*osbH*), the PtdIns(4)*P* 4-phosphatase Sac1, the hydrolase receptor sortilin, the ER proteins calnexin, calmodulin, calreticulin and reticulon, the ER-Golgi intermediate compartment protein-3 (Ergic3), the Golgi proteins golvesin and YIPF1, the vesicle transport through interaction with t-SNAREs homolog 1A (Vti1A), the vesicle-associated membrane protein-7A/B (Vamp7A/B), the vesicle-fusing ATPase (NsfA), the vacuolar protein sorting-associated proteins Vps26, Vps29, and Vps35 (retromer coat complex subunits), dynamin A/B (DymA/B), dynamin-like protein A/C (DlpA/C), Vps4, Vps11, Vps13A, Vps16, Vps45, Vps51, Vps53, syntaxins (5, 7A/B, 8A/B), t-SNAREs, synaptobrevin B, α-SNAP, and many small GTPases or their modulators (Arf1, Arf1 GEF, Arl8, Rab1A/B/C/D, Rab2A/B, Rab4, Rab5A/B, Rab6, Rab7A, Rab8A, Rab11A/C, Rab14, Rab18, Rab21, Rab32A/B/C/D, RabC, RabJ, RabG2, RabQ, Rab GDI, Rac1A, RacB, RacE, RagA, RanA, RanBP1, RasB, RasC, RasG, RasS, RapA, Rap GAP, Rho GAP, Rho GDI, Sar1, Spg1), as well as the interaptin AbpD [67], the metal ion transporter Nramp1 [68, 69], and the GPCR and receptor protein kinase RpkA [70]. Moreover, many mitochondrial proteins were found to associate with LCVs (**Table S1**). Finally, the *L. pneumophila* PtdIns 4-kinase LepB [20], the PtdIns(4)*P*-binding effectors SidC, its paralogue SdcA, and SidM/DrrA [17, 23, 71], as well as the retromer interactor RidL [72] and the deAMPylase SidD [37, 38] were also identified on LCVs from *D. discoideum* Ax3 and Δ*sey1* (**Table S1**).

Furthermore, 74 *D. discoideum* or 34 *L. pneumophila* proteins were identified only on LCVs from *D. discoideum* Ax3, while 11 host and 3 bacterial proteins were present only on LCVs from Δ*sey1* mutant amoeba (**Fig. 1A, Table S1**). Among the host proteins present only on LCVs from *D. discoideum* Ax3, we identified several putative MCS components possibly involved in (PI-driven) lipid transport (OSBP7, Vps13B, CRAL-TRIO domain protein), as well as proteins possibly implicated in ER-Golgi transport (TRAPPC3, YIPF5, γ-SNAP), or localizing to the ER (syntaxin18, calmodulin, Sey1) (**Table S1**). Among the *L. pneumophila* proteins identified only on LCVs from *D. discoideum* Δ*sey1* was the PtdIns(4)*P*-binding effector Lpg2603 [73] (**Table S1**). Based on these findings, we sought to analyze *D. discoideum* factors possibly involved in LCV-ER MCS.

**Figure 1.**
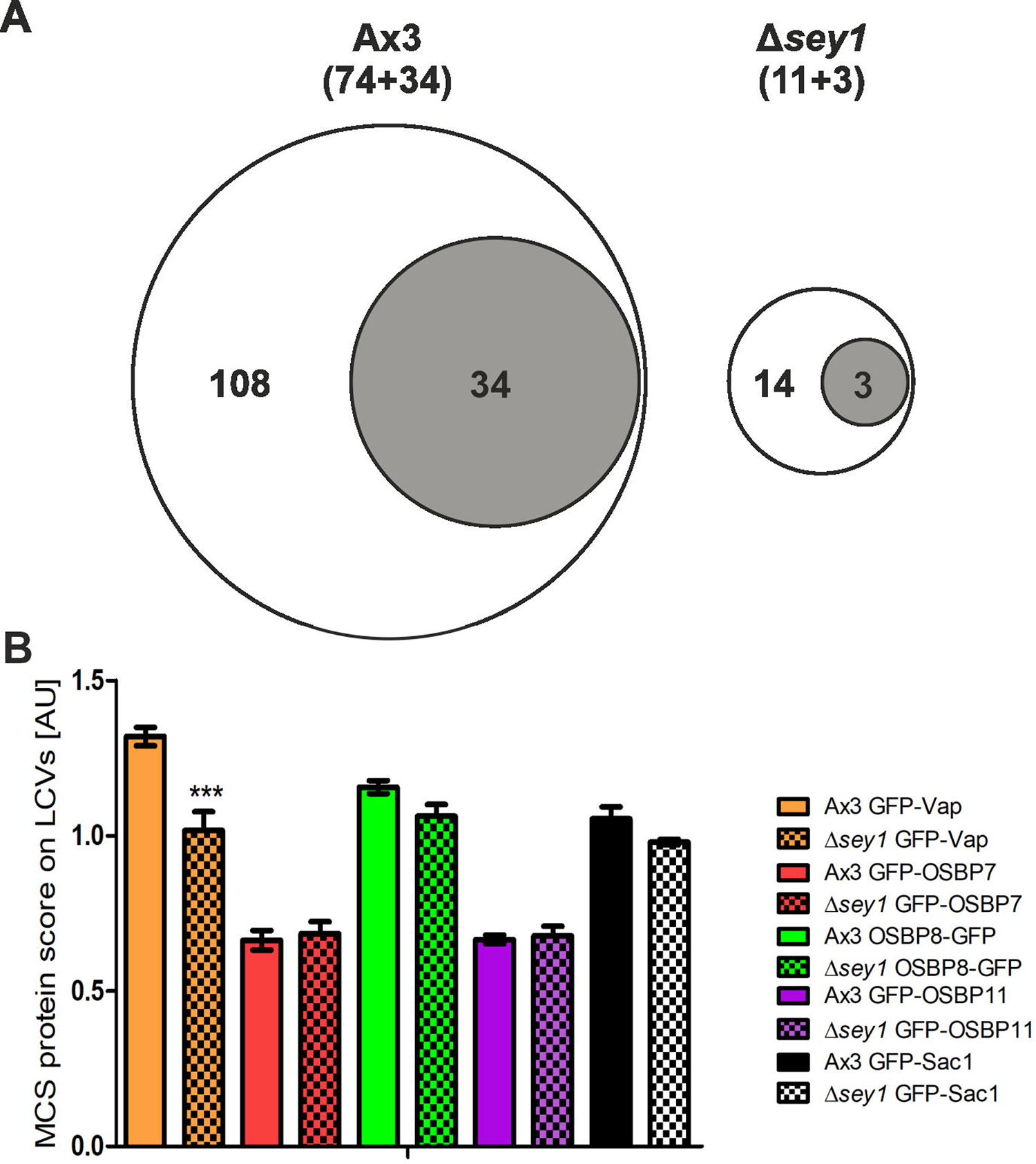
*D. discoideum* MCS components localizing to LCVs. (**A**) Comparative proteomics was performed by mass spectrometry with LCVs purified from *L. pneumophila*-infected *D. discoideum* Ax3 or Δ*sey1* (for details see Materials and Methods). Among the total of 3658 eukaryotic or bacterial proteins identified, the Venn diagrams show proteins identified in biological triplicates only on LCVs from *D. discoideum* Ax3 (108 proteins; 74 *D. discoideum*, 34 *L. pneumophila*) or only on LCVs from Δ*sey1* mutant amoeba (14 proteins; 11 *D. discoideum*, 3 *L. pneumophila*). (**B**) Imaging flow cytometry (IFC) of *D. discoideum* Ax3 or or Δ*sey1* producing P4C-mCherry (pWS032) and GFP fusion proteins of Vap, OSBP7, OSBP8, OSBP11, or Sac1, infected (MOI 5, 1 h) with mPlum-producing *L. pneumophila* JR32 (pAW014). Mean IFC colocalization score and standard error of mean (SEM) for MCS component GFP fusion proteins for > 5000 LCVs is depicted. The data are representative for three independent experiments (***P<0.001).

*D. discoideum* harbors single putative orthologues of the PtdIns(4)*P* 4-phosphatase Sac1 (DDB_G0271630), and the VAMP-associated protein (Vap; DDB_G0278773), as well as several OSBPs, including OSBP7 (DDB_G0283035; *osbG*), OSBP8 (DDB_G0283709; *osbH*) and OSBP11 (DDB_G0288817; *osbK*). All 12 *D. discoideum* OSBPs belong to the class of “short OSBPs”, which harbor a putative lipid-binding ORD domain (OSBP-related domain) but lack ankyrin repeats (protein-protein interaction), a PH domain (PI binding), a trans-membrane domain (TMD; membrane interaction), and the FFAT motif (Vap interaction) (**Fig. S1**). OSBP8 and OSBP11 are most similar to one another and most closely related to human short OSBPs. OSBP7 represents an OSBP more distantly related to the other paralogues, and OSBP1 (*osbA*), OSBP2 (*osbB*), OSBP3 (*osbC*), OSBP5 (*osbE*) and OSBP6 (*osbF*) form a separate cluster.

In order to validate the localization of MCS components to LCVs, GFP fusion proteins of Vap, OSBP7, OSBP8, OSBP11 or Sac1 were co-produced alongside the PtdIns(4)*P*/LCV probe P4C-mCherry [59] in *D. discoideum* Ax3 or Δ*sey1*, and their localization to LCVs was assessed by imaging flow cytometry (IFC) (**Fig. 1B**). All putative MCS components were found to localize to PtdIns(4)*P*-positive LCVs, and Vap accumulated on significantly fewer LCVs in Δ*sey1* mutant amoeba. Since IFC has a rather low spatial resolution, the PtdIns(4)P-positive limiting LCV membrane cannot be discriminated from tightly associated ER. In summary, comparative proteomics identified MCS components on purified LCVs, the localization of which to the pathogen vacuole was confirmed by IFC.

### *D. discoideum* MCS components localize to the ER or the LCV membrane

Based on the above bioinformatic and experimental considerations, we sought to analyze the role of the putative MCS components Vap, OSBP7, OSBP8, OSBP11 and Sac1 for LCV formation and the LCV-ER MCS in detail. Using dually fluorescently labeled *D. discoideum*, we assessed by confocal fluorescence microscopy the co-localization of GFP fusions of Vap, OSBP7, OSBP8, OSBP11 or Sac1 with either the ER-resident protein calnexin A-mCherry (CnxA-mCherry), the PtdIns(4)*P*/LCV probe P4C-mCherry, or the endosomal transporter AmtA-mCherry (**Fig. 2A, Fig. S2A**). This high-resolution approach revealed that Vap and OSBP8, but significantly less OSBP7 and OSBP11, co-localize with CnxA-mCherry in *L. pneumophila*-infected (**Fig. 2B**) as well as in uninfected amoeba (**Fig. S2B-D**). The PtdIns(4)*P* 4-phosphatase Sac1 and its catalytically inactive mutant, Sac1_C383S, also co-localized with CnxA-mCherry, while a Sac1 variant lacking the ER-targeting transmembrane domain, Sac1_ΔTMD, diffusely localized throughout the cell (**Fig. 2B**). Moreover, Vap and OSBP11 co-localized with P4C-mCherry, while OSBP7, OSBP8 and Sac1 did not (**Fig. 2C**), and Vap and OSBP11 also co-localized with AmtA-mCherry more extensively than the other MCS components (**Fig. 2D**). Finally, OSBP11 intensely localized to the PM. In summary, Vap, OSBP8 and Sac1 co-localize with ER-resident CnxA-mCherry, and Vap as well as OSBP11 co-localize with the PtdIns(4)*P* probe P4C-mCherry and the endosomal/LCV marker AmtA-mCherry (**Table 1**). Hence, at LCV-ER MCS Vap localizes to LCVs as well as to the ER, while OSBP8 and Sac1 preferentially localize to the ER, and OSBP11 preferentially localizes to LCVs.

**Figure 2.**
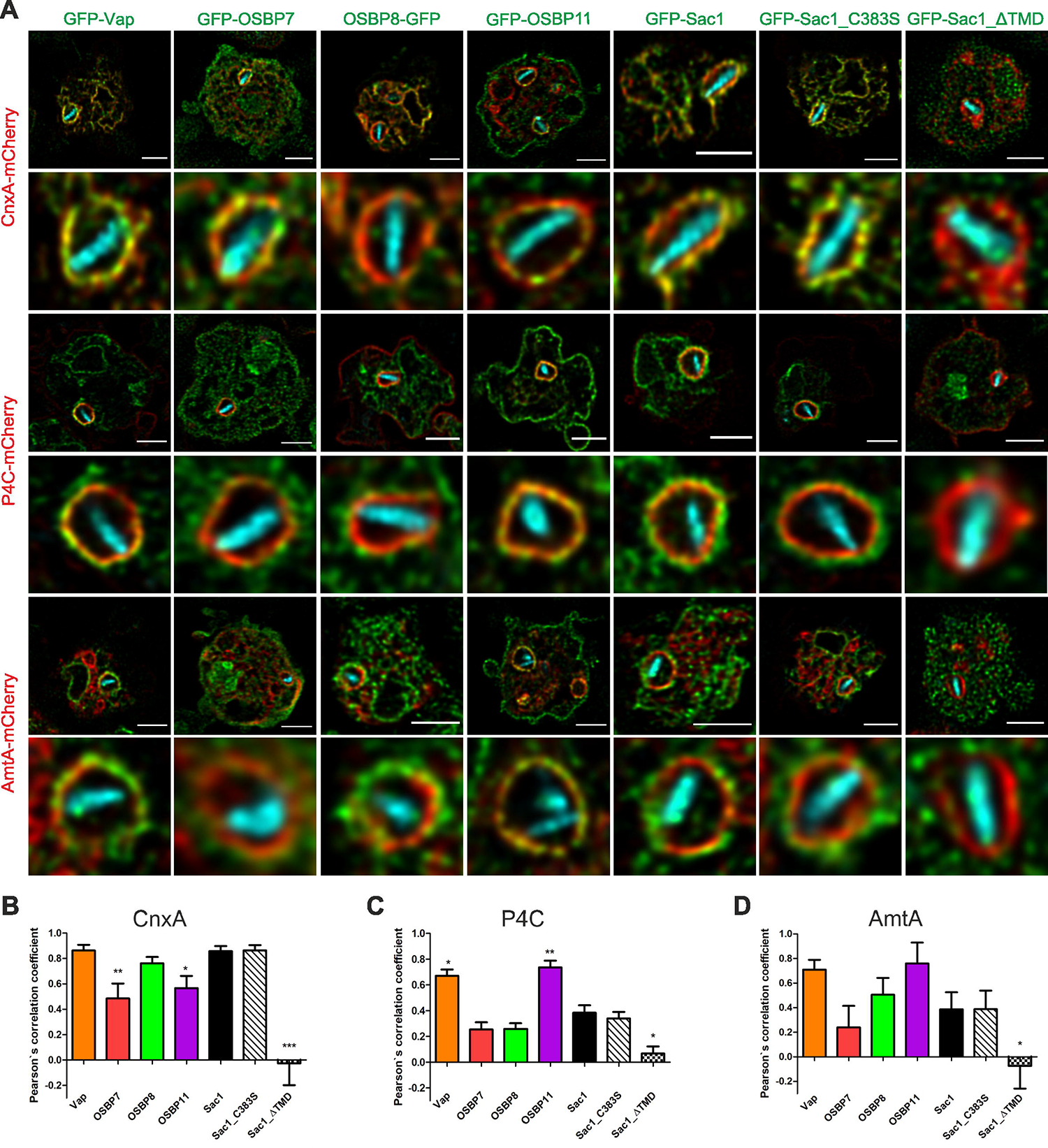
*Dictyostelium discoideum* MCS components localize to LCVs and/or ER. (**A**) Dually labeled *D. discoideum* amoebae producing MCS components fused to GFP, and either calnexin (CnxA)-mCherry (pAW012), P4C-mCherry (pWS032), or AmtA-mCherry were infected (MOI 5, 2 h) with mCerulean-producing *L. pneumophila* JR32 (pNP99), fixed with 4 % PFA, and imaged by confocal fluorescence microscopy. Merged images are shown. Scale bars: 3 μm. The Pearson’s correlation coefficient was generated using Coloc 2 from Fiji (ImageJ) and is shown for MCS components fused to GFP with respect to (**B**) CnxA-mCherry, (**C**) P4C-mCherry, or (**D**) AmtA-mCherry.

**Table 1.** Features of selected *D. discoideum* MCS components.

| MCS component | Colocalization |  |  | *Legionella* Icm<sup>a</sup> |
| --- | --- | --- | --- | --- |
|  | Calnexin<br>(ER)<sup>a</sup> | P4C<br>(PI4P/LCV)<sup>a</sup> | AmtA<br>(PM/EE)<sup>a</sup> |  |
| Vap | ++ | ++ | + | positive |
| OSBP7 (*osbG*) | ++/-<sup>b</sup> | - | - | no effect |
| OSBP8 (*osbH*) | ++ | - | - | negative |
| OSBP11 (*osbK*) | - | ++ | ++ | positive |
| Sac1 | ++ | - | - | no effect |
| Sac1_C383S | ++ | - | - | no effect |
| Sac1_ΔTMD | - | - | - | negative |

<sup>a</sup> Abbreviations: Icm, intracellular multiplication of *L. pneumophila* in *D. discoideum* producing the Sac1 variants or lacking *vap*, *osbG*, *osbH* or *osbK*, compared to replication in the parental strain Ax3; ER, endoplasmic reticulum; EE, early endosome; PI4P/LCV, phosphatidylinositol 4-phosphate/*Legionella*-containing vacuole; PM, plasma membrane.
<sup>b</sup> +: co-localization in *L. pneumophila*-infected cells; -: no co-localization in uninfected cells.

### MCS components modulate the replication of *L. pneumophila* in *D. discoideum* and mammalian cells

To further assess the role of the MCS components for intracellular replication of *L. pneumophila* and LCV formation, we constructed *D. discoideum* deletion mutant strains lacking *vap*, *osbG*, *osbH* or *osbK* by disrupting the corresponding genes with a blasticidin selection marker (**Fig. S3**). Several independent clones of the deletion mutants were picked and further analyzed. Despite repeated attempts, we failed to obtain a Δ*sac1* deletion mutant, and therefore, the gene might be essential in *D. discoideum*. Instead, we constructed and used a dominant negative Sac1 variant lacking the ER-targeting transmembrane domain, Sac1_ΔTMD, and the catalytically inactive Sac1_C383S mutant.

To assess the role of these putative MCS components for intracellular replication of *L. pneumophila*, we used mCherry-producing bacteria and quantified fluorescence intensity (RFU) (**Fig. 3**) and colony forming units (CFU) (**Fig. S4A**). We compared intracellular growth of the wild-type *L. pneumophila* strain JR32 in *D. discoideum* Δ*vap*, as well as in the Δ*osbG*, Δ*osbH*, and Δ*osbK* mutant strains (**Fig. 3A**). In *D. discoideum* Δ*vap*, intracellular bacterial growth was reduced at 4-8 days post infection (p.i.) and enhanced at 10 days p.i. Moreover, intracellular bacterial growth was enhanced in *D. discoideum* Δ*osbH*, reduced in Δ*osbK* and not affected in Δ*osbG*. The intracellular growth phenotypes of *L. pneumophila* in the *D. discoideum* Δ*vap*, Δ*osbH* and Δ*osbK* mutants were reverted by expressing the corresponding genes on a plasmid (**Fig. 3A**). The intracellular growth of *L. pneumophila* strain JR32 was similar in *D. discoideum* Ax3 producing GFP, GFP-Sac1 or GFP-Sac1_C383S but significantly reduced in amoeba producing GFP-Sac1_ΔTMD (**Fig. 3B, Fig. S4A**). These results are in agreement with the notion that Sac1 is implicated in intracellular growth of *L. pneumophila*, and Sac1_ΔTMD but not GFP-Sac1_C383S interfere with growth in a dominant negative manner. An *L. pneumophila* Δ*icmT* mutant, lacking a functional Icm/Dot T4SS, did not replicate in any of the *D. discoideum* mutant strains (data not shown). Taken together, compared to the parental *D. discoideum* strain, *L. pneumophila* replicates less efficiently in absence of the MCS components Vap or OSBP11 and upon the production of Sac1_ΔTMD, while the bacteria replicate more efficiently in absence of OSBP8 (**Table 1**).

**Figure 3.**
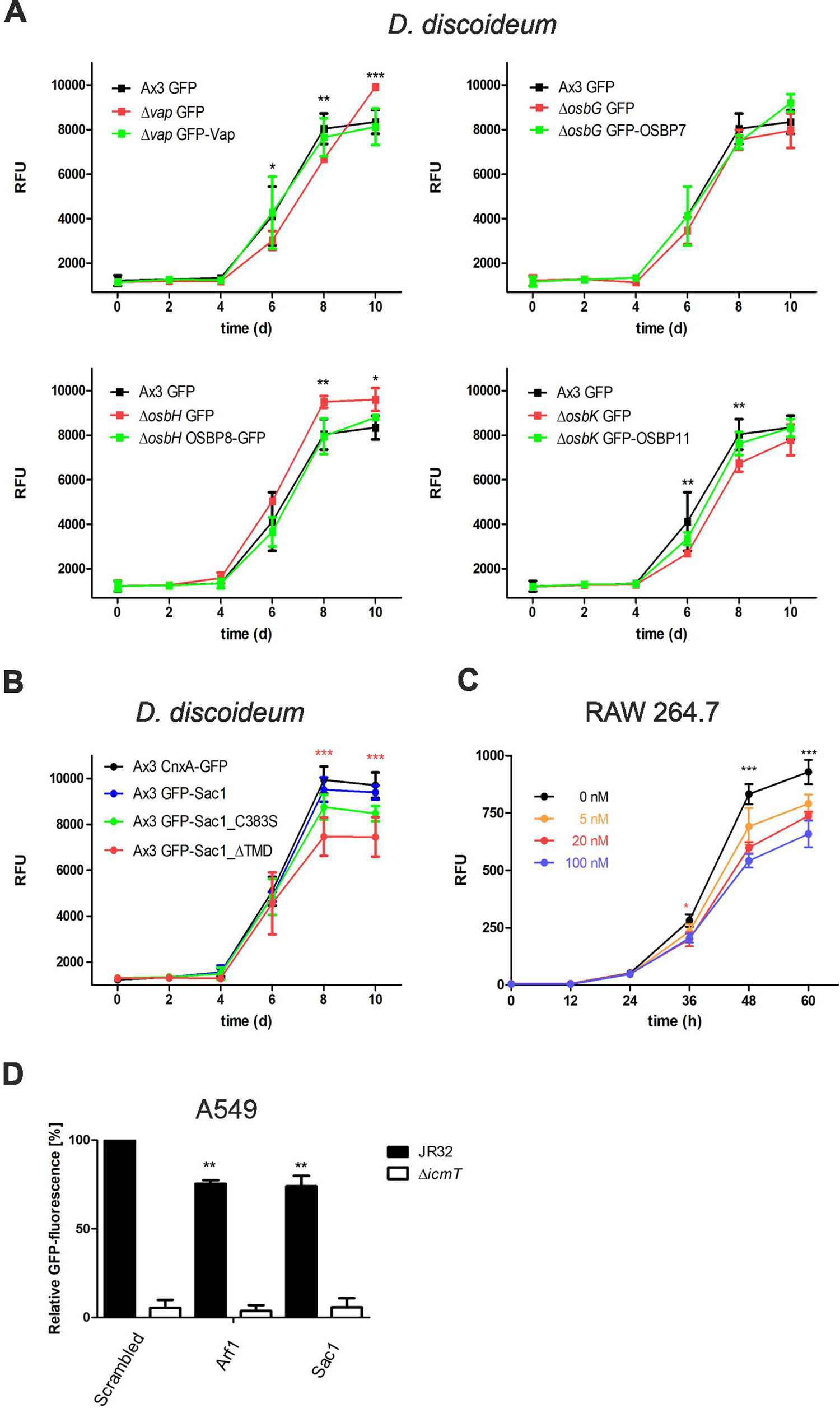
MCS components modulate replication of *Legionella pneumophila* in *Dictyostelium discoideum* and mammalian cells. (**A**) *D. discoideum* Ax3, or the Δ*vap*, Δ*osbG,* Δ*osbH*, or Δ*osbK* mutant strains producing GFP (pDM317) or GFP fusions of the MCS components were infected (MOI 1) with mCherry-producing *L. pneumophila* JR32 (pNP102), and intracellular replication was assessed by relative fluorescence units (RFU). Mean and SEM of three independent experiments are shown (*P<0.05; **P<0.01; ***P<0.001). (**B**) *D. discoideum* Ax3 producing CnxA-GFP (pAW016), GFP-Sac1 (pLS037), GFP-Sac1_C383S (pSV015) or GFP-Sac1_ΔTMD (pSV034) were infected (MOI 1) with mCherry-producing *L. pneumophila* JR32 (pNP102), and intracellular replication was assessed by relative fluorescence units (RFU). Mean and SEM of three independent experiments are shown (***P<0.001). (**C**) RAW 264.7 macrophages were treated with increasing concentrations of OSW-1 (0-100 nM, 1-60 h), infected (MOI 1) with GFP-producing *L. pneumophila* JR32 (pNT28), and intracellular replication was assessed by relative fluorescence units (RFU). Mean and SEM of three independent experiments are shown (*P<0.05; ***P<0.001). (**D**) A549 epithelial cells transfected for 48 h with 10 nM siRNA oligonucleotides were infected (MOI 10) with GFP-producing *L. pneumophila* JR32 or Δ*icmT* (pNT28), and intracellular replication was assessed by fluorescence with a microplate reader. Growth after 24 h was compared to growth at 1 h. Mean and SEM of three independent experiments are shown (**P<0.01).

OSBPL8 was identified on LCVs purified from *L. pneumophila*-infected RAW 264.7 macrophages [55, 56], suggesting that OSBPs and MCS components might also play a role for LCV formation in mammalian cells. To assess the role of MCS components for intracellular replication of *L. pneumophila* in mammalian cells, we used a pharmacological approach and RNA interference. Upon treatment of RAW 264.7 macrophages with the steroidal saponin OSBP inhibitor OSW-1 [65], the intracellular replication of GFP-producing *L. pneumophila* JR32 was significantly inhibited in a dose-dependent manner (**Fig. 3C**). Furthermore, the depletion of Sac1 by RNA interference reduced the intracellular replication of *L. pneumophila* in A549 lung epithelial cells by approximately 25% (**Fig. 3D, Fig. S4BC**), similarly to the depletion of the small GTPase Arf1 used as a positive control [59]. Taken together, pharmacological and RNA interference experiments indicate that MCS components also promote the intracellular replication of *L. pneumophila* in mammalian cells.

### *D. discoideum* Vap, OSBP11 and Sac1 promote the expansion of PtdIns(4)*P*-positive LCVs

Next, we sought to correlate the intracellular replication of *L. pneumophila* in *D. discoideum* lacking MCS components with alterations of the LCVs. To this end, we quantified the LCV area in *D. discoideum* Ax3 and compared it to the LCVs in the Δ*vap*, Δ*osbG*, Δ*osbH*, Δ*osbK* or Δ*sey1* mutants or to LCVs obtained in strain Ax3 upon production of GFP-Sac1_ΔTMD (**Fig. 4A, Fig. S5A**).

**Figure 4.**
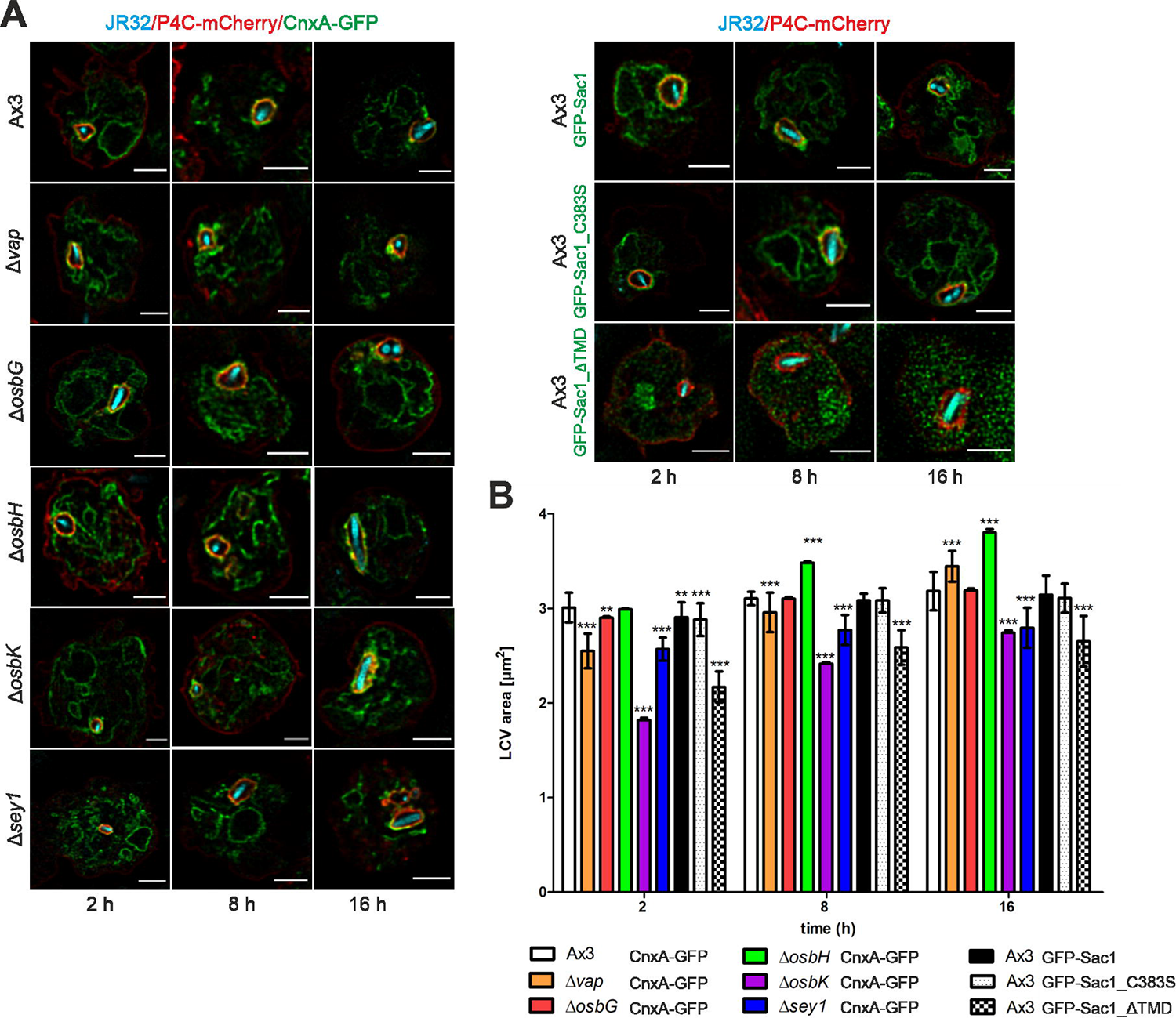
Vap, OSBP11, and Sac1 promote expansion of PtdIns(4)*P*-positive LCVs. (**A**) Dually labeled *D. discoideum* Ax3, Δ*vap*, Δ*osbG,* Δ*osbH*, Δ*osbK* or Δ*sey1* mutants producing P4C-mCherry (pWS032) and CnxA-GFP (pAW016), or Ax3 producing P4C-mCherry and either GFP-Sac1 (pLS037), GFP-Sac1_C383S (pSV015) or GFP-Sac1_ΔTMD (pSV034) were infected (MOI 5, 2-16 h) with mCerulean-producing *L. pneumophila* JR32 (pNP99) and fixed with 4 % PFA. Merged images for the analyzed time points are shown. Scale bars: 3 μm. (**B**) LCV area was measured using ImageJ (n=100-200 per condition from 3 independent biological replicates). Means and SEM of single cells are shown (**P<0.01; ***P<0.001).

Compared to LCVs in *D. discoideum* Ax3, LCVs in the Δ*vap* strain were significantly smaller at 2 h and 8 h p.i., and larger at 16 h p.i. (**Fig. 4B**). Moreover, the LCVs were significantly smaller in the Δ*osbK* mutant strain at 2 h, 8 h and 16 h p.i., larger in the Δ*osbH* mutant at 8 h and 16 h p.i. and did not change in the Δ*osbG* mutant (**Fig. 4B**). Intriguingly, the LCVs in the Δ*osbK* mutant strain were even smaller than in the Δ*sey1* mutant used as a control [66]. The production of GFP-Sac1_ΔTMD in *D. discoideum* Ax3 significantly reduced the LCV size at 2 h, 8 h and 16 h post infection compared to amoeba producing GFP-Sac1 or GFP-Sac1_C383S (**Fig. 4B**). Upon production of GFP-Sac1_ΔTMD in the Δ*vap*, Δ*osbG*, Δ*osbH*, Δ*osbK*, or Δ*sey1* mutant strains the LCV size was also reduced at all time points p.i. compared to amoebae producing GFP-Sac1 (**Fig. S5B**). Taken together, reduced intracellular replication in some *D. discoideum* strains (Δ*vap*, Δ*osbK*, Δ*sey1*, and Sac1_ΔTMD) is positively correlated with a reduced LCV expansion, and increased intracellular replication in the Δ*osbH* mutant is correlated with an enhanced LCV expansion.

### Sac1_**Δ**TMD reduces the accumulation of PtdIns(4)*P*, Vap and OSBP8 on LCV-ER MCS

We then sought to assess the effects of *D. discoideum* MCS components on the LCV PtdIns(4)*P* levels. To this end, we infected amoeba with either mCerulean- or mPlum-producing *L. pneumophila* and analyzed PtdIns(4)*P* on LCVs by confocal microscopy and imaging flow cytometry (IFC) using dually labeled *D. discoideum* strains producing P4C-mCherry and either GFP-Sac1 (**Fig. 5AB, Fig. S6**) or GFP-Sac1_ΔTMD (**Fig. 2A, Fig. 4A, Fig. 5B**, data not shown).

**Figure 5.**
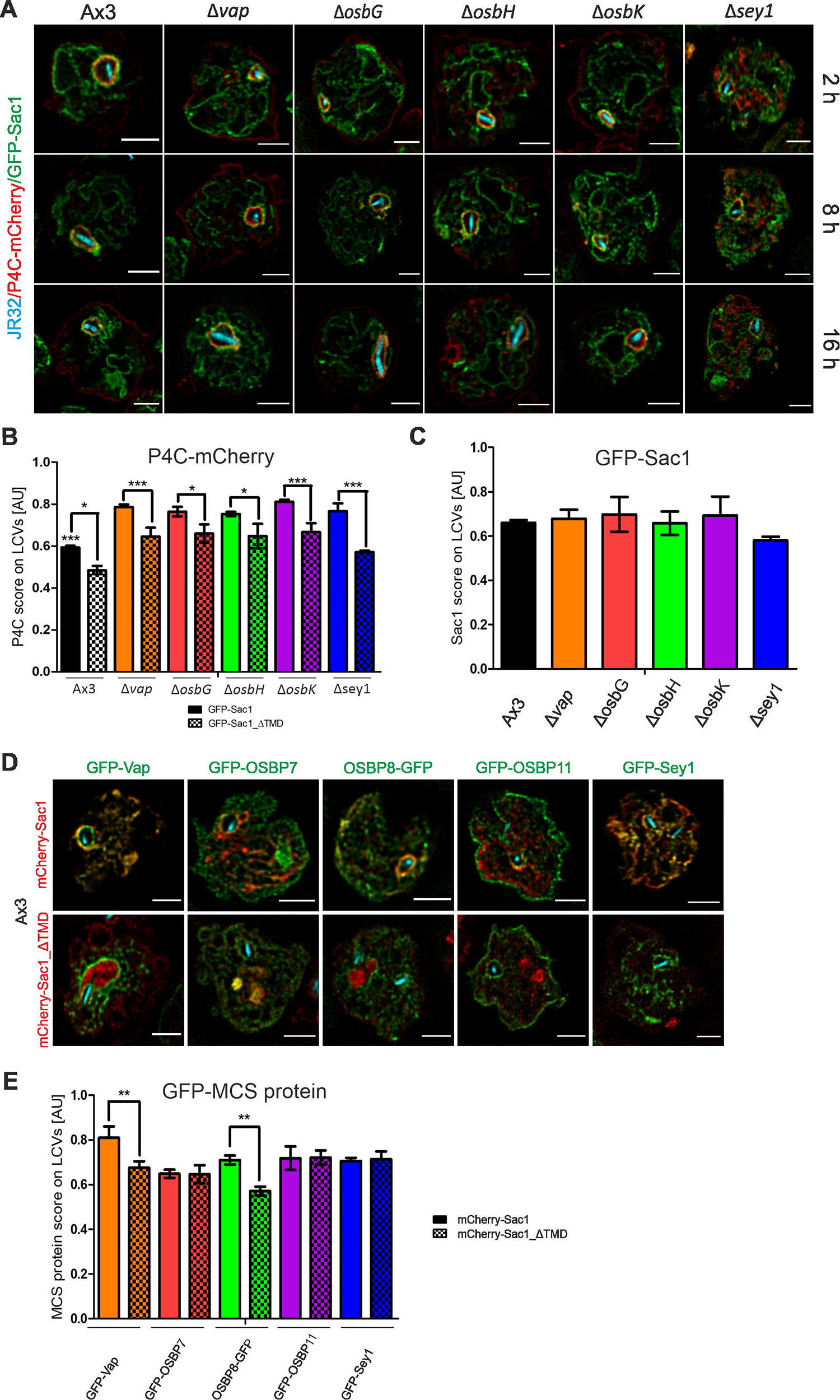
Sac1_ΔTMD reduces the accumulation of PtdIns(4)*P*, Vap and OSBP8 on LCVs. (**A**) Dually labeled *D. discoideum* Ax3, Δ*vap*, Δ*osbG,* Δ*osbH*, Δ*osbK* or Δ*sey1* mutants producing P4C-mCherry (pWS032) and GFP-Sac1 (pLS037) were infected (MOI 5, 2-16 h) with mCerulean-producing *L. pneumophila* JR32 (pNP99) and fixed with 4 % PFA. Merged images for the analyzed time points are shown. Scale bars: 3 μm. (**B, C**) Imaging flow cytometry (IFC) analysis of dually labeled *D. discoideum* Ax3, Δ*vap*, Δ*osbG,* Δ*osbH*, Δ*osbK* or Δ*sey1* mutants producing P4C-mCherry (pWS032) and either GFP-Sac1 (pLS037) or GFP-Sac1_ΔTMD (pSV034), infected (MOI 5, 2 h) with mPlum-producing *L. pneumophila* JR32 (pAW014). (**D**) Dually labeled *D. discoideum* Ax3 producing GFP fusions of MCS components and either mCherry-Sac1 (pSV044) or mCherry-Sac1_ΔTMD (pSV045) were infected (MOI 5, 2 h) with mCerulean-producing *L. pneumophila* JR32 (pNP99) and fixed with 4 % PFA. Merged images for the analyzed time points are shown. Scale bars: 3 μm. (**E**) IFC analysis of dually labeled *D. discoideum* Ax3 producing GFP fusions of MCS components and either mCherry-Sac1 (pSV044) or mCherry-Sac1_ΔTMD (pSV045), infected (MOI 5, 2 h) with mPlum-producing *L. pneumophila* JR32 (pAW014). Quantification of (B) P4C-mCherry (C) Sac1-GFP or (E) GFP fusions of MCS components localizing to LCVs at 2 h p.i. Due to the lower resolution of IFC, “LCVs” designates the LCV limiting membrane and tightly attached ER. Number of events per sample, n = 5000. Data represent mean and SEM of three independent experiments (*P<0.05; **P<0.01; ***P<0.001).

Compared to the ectopic production of GFP-Sac1, the production of GFP-Sac1_ΔTMD in *D. discoideum* Ax3 resulted in a significant decrease of the IFC P4C colocalization score (see Materials and Methods) for the acquisition of P4C-mCherry on LCV-ER MCV (**Fig. 5B**). GFP-Sac1_ΔTMD localized in strain Ax3 throughout the cell and does not appear to have an organelle preference (**Fig. 2A, Fig. 4A**). Accordingly, this result suggests that ectopic production of GFP-Sac1_ΔTMD reduces the PtdIns(4)*P* levels on both the ER as well as the limiting LCV membrane, which cannot be discriminated due to the lower resolution of IFC compared to confocal microscopy. Similarly, the ectopic production of GFP-Sac1_ΔTMD in the *D. discoideum* Δ*vap*, Δ*osbG*, Δ*osbH*, or Δ*osbK* mutant strains resulted in a decrease of PtdIns(4)*P* on LCV-ER MCS (**Fig. 5B**), while GFP-Sac1_ΔTMD localized in the mutant strains throughout the cell (data not shown). At the same time the P4C colocalization score on LCV-ER MCS in the Δ*vap*, Δ*osbG*, Δ*osbH*, or Δ*osbK* mutant strains was similar among the mutants but overall higher than on LCVs in the Ax3 parental strain (**Fig. 5B**). These findings suggest that the depletion of a single MCS component suffices to increase the overall levels of PtdIns(4)*P* on LCV-ER MCS.

Upon production of GFP-Sac1 in the *D. discoideum* Δ*vap*, Δ*osbG*, Δ*osbH*, or Δ*osbK* mutant strains, the IFC Sac1 colocalization score was similar for all LCV-ER MCS (**Fig. 5C**). This result indicates that the localization of Sac1 to LCV-ER MCS is not dependent on Vap or the OSBPs tested. The Sac1 colocalization score was slightly (but not significantly) lower in the Δ*sey1* mutant strain, suggesting that Sey1 might play a role in the acquisition of Sac1 to LCV-ER MCS (**Fig. 5C**). This result is reflected in the finding that Sey1 promotes the recruitment of ER to LCVs [66]. In summary, compared to the production of GFP-Sac1, the production of GFP-Sac1_ΔTMD in *D. discoideum* Ax3 or the Δ*vap*, Δ*osbG*, Δ*osbH*, Δ*osbK* or Δ*sey1* mutant strains decreased the PtdIns(4)*P* level on LCV-ER MCS, while Sac1 did not change, in agreement with the notion that GFP-Sac1_ΔTMD globally reduces PtdIns(4)*P* levels.

We also assessed by confocal microscopy and IFC the effect of Sac1_ΔTMD for the accumulation of Vap, OSBP7, OSBP8 or OSBP11 on LCVs (**Fig. 5DE, Fig. S7**). To this end, we used dually fluorescent labeled *D. discoideum* Ax3 producing either mCherry-Sac1 or mCherry-Sac1_ΔTMD and GFP fusion proteins of Vap, OSBP7, OSBP8 or OSBP11. Compared to the ectopic production of mCherry-Sac1, the production of mCherry-Sac1_ΔTMD in *D. discoideum* Ax3 resulted in a significant decrease of the IFC MCS component colocalization score for the acquisition on LCV-ER MCS of GFP fusions of Vap and OSBP8, but not OSBP7, OSBP11 or Sey1 (**Fig. 5E**). Taken together, compared to the production of mCherry-Sac1, the production of mCherry-Sac1_ΔTMD in *D. discoideum* Ax3, decreased the levels of Vap and OSBP8 on LCV-ER MCS, suggesting that the localization of these MCS components is regulated by PtdIns(4)*P* levels on the LCVs.

### The *L. pneumophila* effectors LepB and SidC promote expansion of PtdIns(4)*P*-positive LCVs

To assess the role of *L. pneumophila* effector proteins for LCV remodeling and PtdIns(4)*P* levels, we analyzed LepB, a Rab1 GTPase activating protein (GAP)/PtdIns 4-kinase [20, 31] and SidC, a PtdIns(4)*P* interactor/ubiquitin ligase [17, 57, 74, 75]. To this end, we quantified the LCV area in *D. discoideum* Ax3 producing CnxA-GFP, GFP-Sac1 or GFP-Sac1_ΔTMD, following an infection with mCerulean-producing *L. pneumophila* JR32, Δ*lepB* or Δ*sidC* (**Fig. 6A, Fig. S8**).

**Figure 6.**
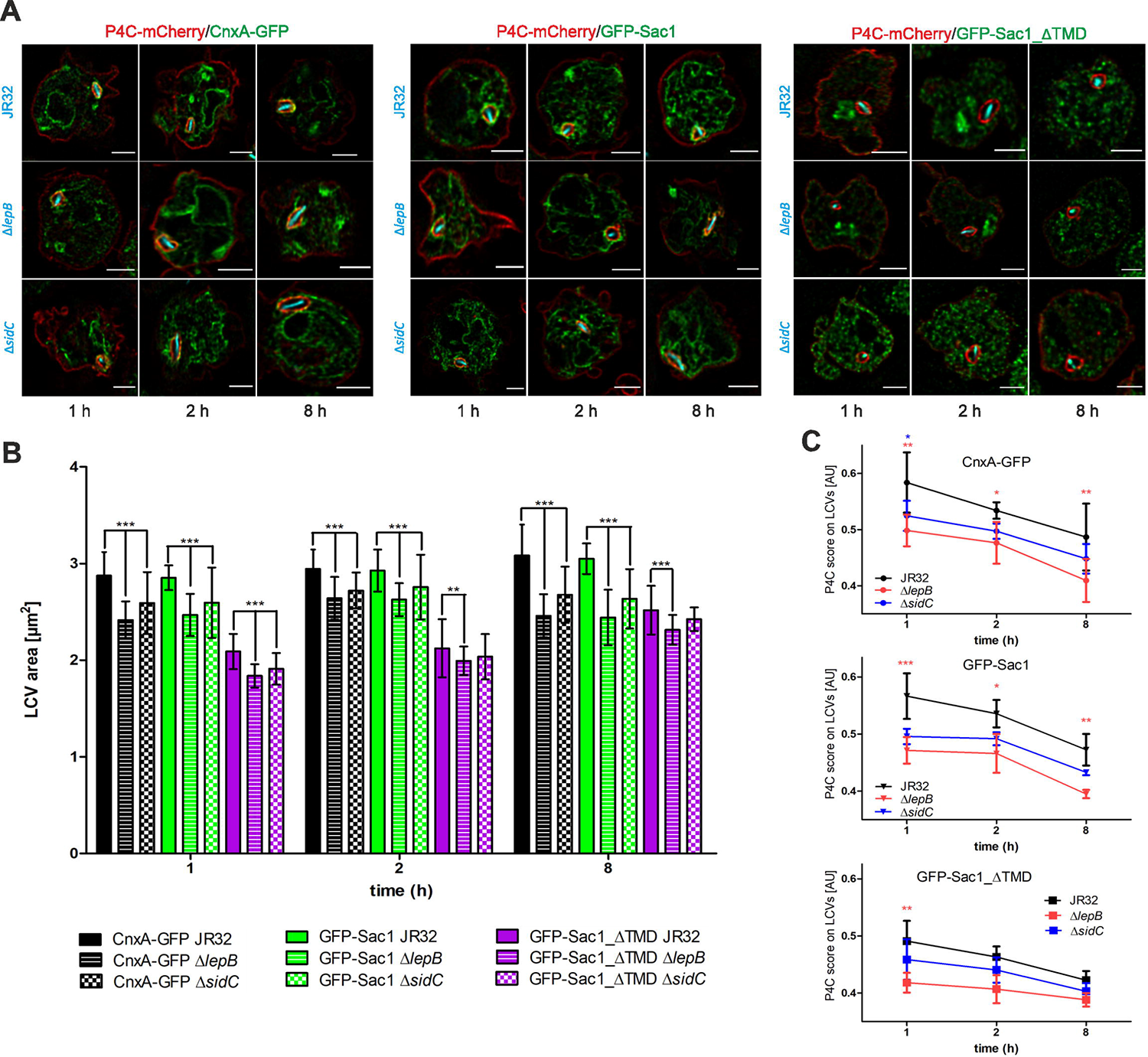
The *L. pneumophila* effectors LepB and SidC determine PtdIns(4)*P* decoration and expansion of LCVs. (**A**) Dually labeled *D. discoideum* Ax3 producing P4C-mCherry (pWS032) and either CnxA-GFP (pAW016), GFP-Sac1 (pLS037), or GFP-Sac1_ΔTMD (pSV034) were infected (MOI 5, 1-8 h) with mCerulean-producing *L. pneumophila* JR32, Δ*lepB* or Δ*sidC* (pNP99) and fixed with 4 % PFA. Merged images for the analyzed time points are shown. Scale bars: 3 μm. (**B**) LCV area was measured using ImageJ (n=100-200 per condition). Means and SEM of single cells are shown (**P<0.01; ***P<0.001). (**C**) Imaging flow cytometry (IFC) analysis of dually labeled *D. discoideum* Ax3 producing P4C-mCherry (pWS032) and either CnxA-GFP (pAW016), GFP-Sac1 (pLS037), or GFP-Sac1_ΔTMD (pSV034), infected (MOI 5, 1-8 h) with mPlum-producing *L. pneumophila* JR32 (pAW014). Number of events per sample, n = 5000. Data represent mean and SEM of three independent experiments (*P<0.05; **P<0.01; ***P<0.001).

In *D. discoideum* producing CnxA-GFP, LCVs harboring Δ*lepB* or Δ*sidC* mutant *L. pneumophila* were significantly smaller than LCVs harboring strain JR32 at 1 h, 2 h and 8 h p.i. (**Fig. 6B**). Overall, the LCVs were of similar size in *D. discoideum* producing GFP-Sac1, but significantly smaller in *D. discoideum* producing GFP-Sac1_ΔTMD. In the latter case, LCVs harboring Δ*lepB* or Δ*sidC* mutant *L. pneumophila* were also significantly smaller than LCVs harboring strain JR32 at 1 h, 2 h and 8 h p.i. (Δ*lepB*) and at 1 h p.i. (Δ*sidC*).

In order to correlate the LCV area with the PtdIns(4)*P* levels on the pathogen compartment, we used IFC and *D. discoideum* strains producing in parallel P4C-mCherry and CnxA-GFP, GFP-Sac1 or GFP-Sac1_ΔTMD, infected with mPlum-producing *L. pneumophila* JR32, Δ*lepB* or Δ*sidC* (**Fig. 6C**). In all cases, the IFC colocalization score for the acquisition of P4C-mCherry on LCVs decreased in the course of an infection (1-8 h p.i.). Compared to the parental strain JR32, the score was lowest for Δ*lepB* and intermediate for Δ*sidC*. The colocalization scores were similar in *D. discoideum* producing CnxA-GFP or GFP-Sac1, but significantly lower in *D. discoideum* producing GFP-Sac1_ΔTMD. Taken together, the lack of the *L. pneumophila* Rab1 GAP/PtdIns 4-kinase LepB or the PtdIns(4)*P* interactor/ubiquitin ligase SidC causes a reduction in LCV area and PtdIns(4)*P* levels. These results indicate that the *L. pneumophila* effector proteins LepB and SidC play a role in the PtdIns(4)*P*-dependent LCV remodeling at LCV-ER MCS.

## Discussion

In this study, we identified several *D. discoideum* MCS components localizing to LCVs (**Fig. 1**). We then used dually fluorescence-labeled *D. discoideum* amoeba to visualize and quantify MCS components at the LCV-ER interface. The protein Vap localized to the PtdIns(4)*P*-positive LCV membrane as well as the ER, while OSBP8 and Sac1 exclusively localized to the ER, and OSBP11 was detected solely on the LCV membrane (**Fig. 2, Table 1**). The MCS components were found to be implicated in intracellular growth of *L. pneumophila* (**Fig. 3**) and LCV remodeling (**Fig. 4**). Host and bacterial factors regulate the PtdIns(4)*P* levels on LCVs: a derivative of the PtdIns(4)*P* 4-phosphatase Sac1 lacking its membrane anchor, Sac1_ΔTMD (**Fig. 5**), or the lack of the *L. pneumophila* PtdIns 4-kinase, LepB (**Fig. 6**), reduced the PtdIns(4)*P* score on LCVs. Taken together, these findings are compatible with the notion that a *Legionella*- and host cell-driven PtdIns(4)*P* gradient at LCV-ER MCSs promotes Vap-, Sac1- and OSBP-dependent pathogen vacuole remodeling (**Fig. 7**).

**Figure 7.**
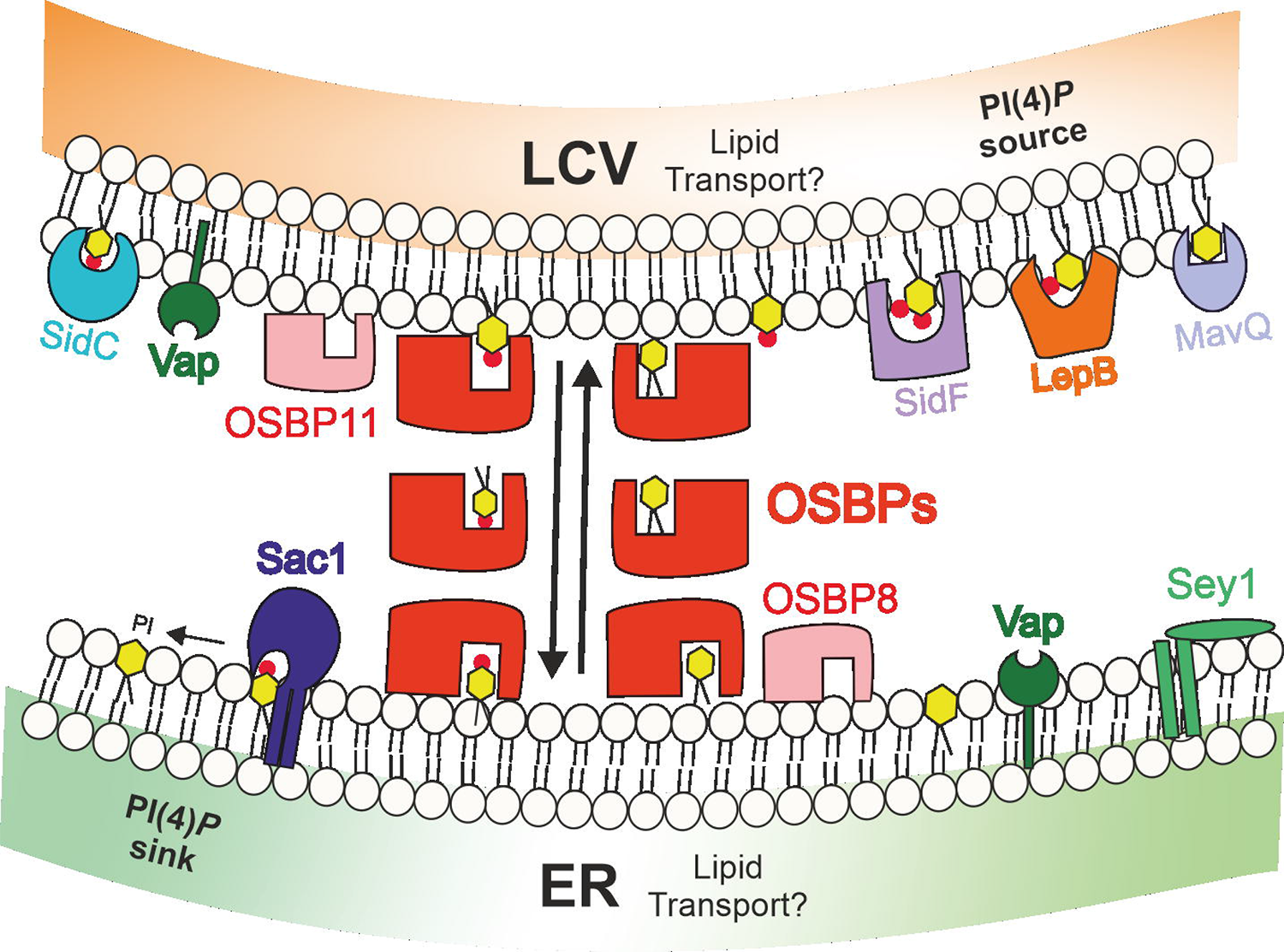
Localization and function of host and bacterial factors at LCV-ER MCS. In dually fluorescence-labeled *D. discoideum* amoeba MCS components such as Vap, OSBPs and Sac1 localize to the LCV-ER interface. The host proteins Vap, OSBP8, Sac1 and Sey1 localize to the ER, and OSBP11 is detected on the PtdIns(4)*P*-positive LCV membrane as well as on the PM. The MCS components are implicated in intracellular growth of *L. pneumophila* and LCV remodeling. The PtdIns(4)*P* 4-phosphatase Sac1 lacking its membrane anchor, Sac1_ΔTMD, or the lack of the *L. pneumophila* PtdIns(3)*P* 4-kinase, LepB, reduced the PtdIns(4)*P* score on LCV-ER MCS. These findings suggest that a *Legionella*- and host cell-driven PtdIns(4)*P* gradient at LCV-ER MCS promotes Vap-, OSBP- and Sac1-dependent pathogen vacuole remodeling.

We quantified LCV size changes at early infection time points (1-2 h p.i.) to assess the role of MCS components in LCV remodeling (**Fig. 4, Fig. 6**). This approach revealed that Vap and even more pronouncedly OSBP11 promote LCV expansion. The size reduction of LCVs in *D. discoideum* Δ*vap* or Δ*osbK* mutant strains was relatively small: the LCVs in the mutant strains were ca. 2-2.5 µm^2^, compared to ca. 3 µm^2^ in the Ax3 parental strain. Accordingly, these small LCV size changes likely reflect a structural remodeling of the pathogen vacuole rather than a substantial LCV expansion. MCS promote the non-vesicular exchange of lipids in mammalian cells [60], and “counter lipids” transported by OSBPs along a Sac1-dependent PtdIns(4)*P* gradient at ER MCS include phosphatidylserine [63, 64] and cholesterol [61, 65]. The identification of the lipids exchanged by the LCV-ER PtdIns(4)*P* gradient and the specific OSBPs involved will be the subject of future studies.

A massive expansion of the pathogen vacuole at later infection time points (> 8 h p.i.) is required to accommodate intracellular replication of *L. pneumophila* and is likely promoted by vesicle fusion. Indeed, LCV formation implicates the interception of (anterograde and retrograde) vesicular trafficking between the ER and the Golgi apparatus [42–44], and nascent LCVs continuously capture and accumulate PtdIns(4)*P*-positive vesicles from the Golgi apparatus [49]. Further supporting the notion that ER-derived vesicles fuse with LCVs is the finding that the SNARE Sec22b, which promotes fusion of ER-derived vesicles with the *cis*-Golgi, is recruited to LCVs shortly after infection [45, 46]. The fusion requires the *L. pneumophila* effector-mediated activation of Rab1 on LCVs and is mediated by the non-canonical pairing of the v-SNARE Sec22b on the vesicles with PM-derived t-SNAREs such as syntaxin2, −3, −4 and SNAP23 on the LCV [47, 48]. In agreement with this scenario, syntaxins (5, 7A/B, 8A/B, 18), t-SNAREs, α- and γ-SNAP as well as Vti1A, Vamp7A/B and NsfA were identified on LCVs by proteomics (**Table S1**).

Moreover, several Icm/Dot-translocated *L. pneumophila* effector proteins were identified on LCVs by the comparative proteomics approach (**Table S1**). These effectors prominently include several PtdIns(4)*P*-binding *L. pneumophila* enzymes: the ubiquitin ligase SidC and its paralogue SdcA, the Rab1 GEF/AMPylase SidM/DrrA and the phytate-activated protein kinase Lpg2603. Furthermore, the deAMPylase and SidM antagonist SidD, as well as the retromer interactor RidL were identified on purified LCVs. To study the role of effector proteins for LCV remodeling and PtdIns(4)*P* levels, we analyzed the Δ*lepB* and Δ*sidC* mutant strains (**Fig. 6**). LCVs harboring these mutant strains were smaller and showed a reduced P4C score at LCV-ER MCS. Noteworthy, the effects were augmented in *D. discoideum* producing Sac1_ΔTMD. These results are readily explicable for the PtdIns 4-kinase LepB: if this kinase is lacking, the PtdIns(4)*P* levels on LCVs are lower and LCV remodeling is impaired. The findings are more difficult to interpret for SidC. This effector localizes exclusively on LCVs in *L. pneumophila*-infected *D. discoideum* [54–57] and tightly binds PtdIns(4)*P* with a dissociation constant, K_d_, in the range of 240 nM [76]. Hence, SidC might titrate PtdIns(4)*P* levels on LCVs, and its absence in *L. pneumophila* is expected to expose more PtdIns(4)*P* on the pathogen vacuole. Accordingly, the finding that LCVs harboring the Δ*sidC* mutant strain show a lower P4C score is caused by more indirect features of the effector.

In addition to the ER-localizing PtdIns(4)*P* 4-phosphatase Sac1, other host cell factors involved in PtdIns(4)*P* turnover might play a role at LCV-ER MCS. The PM-derived PtdIns 4-kinase IIIα (PI4KIIIα) [22] and trans-Golgi-derived PI4KIIIβ [23] promote the membrane localization of PtdIns(4)*P*-binding effectors, and therefore, might modulate PtdIns(4)*P* levels on LCVs. Moreover, while the PtdIns(4,5)*P*_2_ 5-phosphatase OCRL/Dd5P4 produces PtdIns(4)*P* on LCVs, it restricts intracellular replication of *L. pneumophila* [24, 72], suggesting that this PI phosphatase exerts pleiotropic and complex functions in the context of LCV formation and intracellular bacterial replication.

The association of the LCV with ER in amoeba and macrophages represents a longstanding observation [51, 52], which is not understood on a molecular level. In particular, the identity of the tiny, periodic “hair-like” structures between the pathogen vacuole and the ER is unknown. On a similar note, the identity of possible (bacterial or host) tether proteins is not known. A possible candidate for a *L. pneumophila* tether protein is the Icm/Dot-secreted effector Ceg9. This effector might stabilize the LCV-ER contact sites by directly binding to the host protein Rtn4 [58].

Several intracellular bacterial pathogens hijack MCS components during infection [77]. The mechanistically best characterized case is *Chlamydia trachomatis*, which forms a replication-permissive compartment termed inclusion [77]. Intriguingly, the *Chlamydia* integral membrane proteins IncV and IncD tether the inclusion to the ER [78]. While IncV directly binds Vap on the ER through a FFAT motif [79], IncD indirectly makes contact to Vap through the FFAT motif-containing host protein CERT (ceramide transfer protein) [80, 81].

Taken together, this study has revealed the importance of host and bacterial factors for MCS formation, LCV remodeling and intracellular replication of *L. pneumophila*. The study paves the way for an in-depth functional and structural analysis of LCV-ER MCS, and future studies will identify possible *L. pneumophila* tethering factors, which promote and stabilize the LCV-ER MCS.

## Materials and Methods

### Molecular cloning

All plasmids constructed and used are listed in the Supplementary Information (**Table S2**). Cloning was performed using standard protocols, plasmids were isolated by using commercially available kits from Macherey[Nagel, DNA fragments were amplified using Phusion High Fidelity DNA polymerase. For Gibson assembly, the NEBuilder HiFi DNA assembly kit was used. All constructs were verified by DNA sequencing.

To construct the knockout vectors for *vap*, *osbG*, and *osbK* (**Table S2**), the corresponding 5’ fragments were amplified from *D. discoideum* genomic DNA with specific primer pairs oL1277/oL1278, oL1256/oL1356, or oL1687/oL1688, respectively, and cloned into the pBlueScript vector (Stratagene, La Jolla, CA). The 3’ fragments were identically produced using the matching primer pairs, oL1279/oL1280, oL1358/oL1359, or oL1693/oL1694, respectively and cloned into pBlueScript containing the 5’ fragment. After plasmid sequence verification, the blasticidin resistance cassette was inserted between the two 5’and 3’ fragments. The final knockout vectors were digested with KpnI and NotI restriction enzymes before electroporation into Ax3 cells. Blasticidin selection (10 μg/ml) was applied 24 h after transfection. Individual colonies were tested by PCR to confirm gene replacement (**Fig. S3**). The knockout vectors for *osbH* (#629), a kind gift of Prof. Markus Maniak (University of Kassel, Germany), was digested by SpeI and SphI before transfection of Ax3 cells, and transfectants were selected as described above.

The *lepB* deletion strain (IH03) was generated essentially as described [83] by double homologous recombination allelic exchange using counter-selection on sucrose (Suc). The 3’ and 5’ flanking regions of lepB (*lpg2490*) were amplified by PCR using the primer pairs oIH011/oIH013 and oIH012/oIH014, respectively, and genomic *L. pneumophila* DNA as a template. The flanking regions, the Kan^R^ cassette and the pLAW344 backbone cut with the appropriate restriction enzymes were assembled using a four-way ligation, yielding the suicide plasmid pIH29. *L. pneumophila* JR32 was transformed by electroporation with pIH29, and co-integration of the plasmid was assayed by selection on CYE/Km (5-7 days, 30°C). Several clones thus obtained were picked and re-streaked on CYE/Km, grown overnight in AYE medium (96-well plates, 180 rpm) and streaked on CYE/Km/2% Suc. After 3-5 days at 37°C, single colonies were spotted on CYE/Cm, CYE/Km/2% Suc and CYE/Km plates to screen for Cm^S^, Km^R^, Suc^R^ colonies. Double-cross-over events (deletion mutants) were confirmed by PCR screening and sequencing.

The constructs pMIB39 (*gfp*-*osbK*), pMIB41 (*gfp*-*vap*), pMIB87 (*gfp*-*osbG*), and pMIB89 (*osbH*-*gfp*) were constructed by PCR amplification using the primers oMIB38/oMIB39, oMIB14/oMIB12, oMIB20/oMIB18 or oMIB21/oMIB22 and cloned into plasmid pDM317 (*osbK*, *vap*, *osbG*) or pDM323 (*osbH*) after digestion with XhoI/SpeI (*osbK*), BamHI/SpeI (*vap*) or BglII/SpeI (*osbG*, *osbH*).

A GFP-Sac1 fusion was constructed by PCR amplification of the putative open reading frame using *D. discoideum* genomic DNA and the primers oLS039 and oLS040 respectively. The DNA fragment was cloned into BglII and SpeI sites of pDM317, yielding pLS037 (GFP-Sac1). The GFP-Sac1 catalytically inactive mutant (Sac1_C383S) was obtained exchanging the codon TGT (cysteine) in position 383 to AGC coding for serine. Nucleotide substitution was carried out by site-directed mutagenesis according to the manufacturer’s recommendation (QuickChange, Agilent) using pLS037 as template and the PAGE purified primers oSV159 and oSV161 yielding pSV015 (GFP-Sac1_C383S). The truncated GFP-Sac1_ΔTMD was obtained by deletion of the transmembrane domain of the *D. discoideum sac1* gene. To this end, pLS037 excluding the transmembrane domain was amplified with the primers oSV227 and oSV228 yielding pSV034. The plasmids pSV044 and pSV045 harboring Sac1 or Sac1_ΔTMD fused to mCherry were constructed by cloning the genes into BglII and SpeI sites of pDM1042. pLS037 and pSV034 were used as templates.

### Bacteria, cells, growth conditions, and transformation

Bacterial strains and cell lines used are listed in **Table S2**. *L. pneumophila* strains were grown 3 days on charcoal yeast extract (CYE) agar plates, buffered with *N*-(2-acetamido)-2-aminoethane sulfonic acid (ACES) at 37°C. Liquid cultures in ACES yeast extract (AYE) medium were inoculated at an OD_600_ of 0.1 and grown at 37°C for 21 h to an early stationary phase (2 x 10^9^ bacteria/ml). Chloramphenicol (5 µg/ml) was added as required.

*D. discoideum* stains were grown at 23°C in HL5 medium (ForMedium). Transformation of axenic *D. discoideum* amoeba were performed as described [18, 82]. Blasticidin (10 μg/mL), geneticin (G418, 20 µg/ml) and hygromycin (50 µg/ml) were appropriately added.

Murine macrophage-like RAW 264.7 cells and human A549 lung epithelial carcinoma cells were cultivated in RPMI 1640 medium (Life Technologies) supplemented with 10% heat-inactivated fetal bovine serum (FBS: Life Technologies) and 1% glutamine (Life Technologies). The cells were incubated at 37°C with 5% CO_2_ in a humidified atmosphere.

### Intracellular replication of *Legionella pneumophila*

Intracellular growth of *L. pneumophila* JR32 and Δ*icmT* in different *D. discoideum* strains was assessed by measuring fluorescence increase during intracellular replication of mCherry[producing *Legionella* strains (relative fluorescence units, RFU) and by determining colony[forming units (CFU).

To determine intracellular replication of mCherry-producing *L. pneumophila*, *D. discoideum* amoebae were seeded (2 × 10^4^ cells) in 96-well culture-treated plates (Thermo Fisher) and cultured in HL-5 medium at 23°C. The cells were infected (MOI 1) with the *L. pneumophila* strains JR32 or Δ*icmT* harboring plasmid pNP102. After 21 h growth in AYE medium, the bacteria were diluted in MB medium, centrifuged (450 × *g*, 10 min, RT), and incubated for 1 h at 25°C (well plate was kept moist with water in extra wells). Subsequently, infected cells were washed three times with MB medium, incubated for the time indicated at 25°C, and mCherry fluorescence was measured every 2 days using a microtiter plate reader (Synergy H1, Biotek).

To assess CFUs, *D. discoideum* amoebae were infected (MOI 1) with mCherry-producing *L. pneumophila* JR32 or Δ*icmT* grown for 21 h, diluted in MB medium, centrifuged and incubated for 1 h at 25°C. The infected cells were washed three times with MB medium and incubated for the time indicated at 25°C. The infected amoebae were lysed for 10 min with 0.8% saponin (47036, Sigma-Aldrich). Subsequently, serial dilutions were plated every 2 days on CYE agar plated containing Cam (5 μg/ml) and incubated for 3 days at 37°C. CFUs were counted using an automated colony counter (CounterMat Flash 4000, IUL Instruments, CounterMat software) and the number of CFUs (per ml) was calculated.

To test if members of the OSBP family affect intracellular replication of *L. pneumophila* in murine RAW 264.7 macrophages, the steroidal saponin OSBP inhibitor OSW-1 (CAY-30310, Cayman) was added at different concentrations. The macrophages were detached diluted with pre-warmed supplemented RPMI to a concentration of 2.5 × 10^4^ cells/140 μL medium and grown for 24 h in 96-well plates. Liquid cultures of GFP-producing *L. pneumophila* JR32 (pNT28) were grown for 21 h and diluted in pre-warmed supplemented RPMI to a concentration of 5 × 10^4^ cells/50 μL medium. A 10 μM OSW-1 stock solution was prepared freshly. Bacterial suspensions, the OSW-1 working solution (final concentration 5, 20 or 100 nM) and sterile DPBS were added to wells, yielding an MOI 1. The infection was synchronized by centrifugation (450 × *g*, 10 min.). Plates were incubated at 37°C and 5 % CO_2_ in a humidified atmosphere. At the indicated time points (12, 24, 36, 48, 60 h p.i.) intracellular replication was assessed by measuring bacterial GFP production with a plate reader.

### RNA interference, determination of proteins depletion efficiency and cytotoxicity

RNA interference experiments were performed as described [59]. Briefly, A549 cells were treated for 48 h with siRNA oligonucleotides with a final concentration of 10 nM in a 96-well plate (RFU) or 24-well plate (toxicity, Western blot) (**Table S3**). The diluted siRNA was added to the wells (5-10 min) at room temperature (RT), and diluted cells (96-well: 2 × 10^4^, 24-well: 6 × 10^4^) in RPMI medium with FBS was added on top (48 h). Subsequently, the cells were infected (MOI 10) with GFP-producing *L. pneumophila* strains (pNT28), centrifuged and incubated for 1 h at 37°C with 5% CO_2_, washed three times and incubated for 24 h. GFP fluorescence was measured as described above at 1 h and 24 h post infection.

Protein depletion efficiency was assessed as follows: cells were harvested in ice[cold PBS and lysed with ice[cold NP[40 cell lysis buffer. Cell extracts were subjected to SDS PAGE. After Western blotting, PVDF membranes were blocked with PBS/5% bovine serum albumin (BSA; Sigma[Aldrich) for 1 h at RT. Subsequently, specific primary antibodies against SacM1L (13033-1-AP, Proteintech) or GAPDH (2118, Cell Signaling) were diluted 1:500 - 1:1,000 in blocking buffer and used to stain the indicated proteins (4°C, overnight). Finally, horse radish peroxidase (HRP)[conjugated secondary antibodies (GE Healthcare Life Sciences) were diluted 1:2,000 in blocking buffer and incubated (1 h, RT). After extensive washing, the enhanced chemiluminescence (ECL) signal was detected with an ImageQuant LAS4000 (GE Healthcare Life Sciences).

To assess cell viability after siRNA treatment, the Zombie Aqua fixable viability kit (BioLegend) was used. A549 cells were grown and treated with siRNA oligonucleotides (**Table S3**) as described above (protein depletion efficiency). Cells treated for 1 h with 70% sterile[filtered ethanol (EtOH) served as positive control for cell death. The cells were then harvested in ice[cold PBS and stained for 30 min in the dark with 50 µl Zombie Aqua dye 1:500 diluted in PBS. Cells were washed once with supplemented RPMI, centrifuged, washed once with PBS, centrifuged and fixed with 4 % paraformaldehyd (PFA) for 30 min at RT. After centrifugation cells were resuspended in 500 µl PBS. Subsequently, cells were subjected to flow cytometry analysis (BD FACS Canto II). Gates were set according to forward/sideward scatter properties, and 10,000 events were collected for each sample. Mean and standard error of mean (SEM) of Zombie[positive cells are shown.

### Confocal fluorescence microscopy of infected *D. discoideum*

Dually fluorescence labeled *D. discoideum* strains were infected, fixed and imaged by confocal fluorescence microscopy. Prior to infection, exponential phase *D. discoideum* amoebae were seeded (1 × 10^5^ cells per well) in HL5 medium containing geneticin (G418, 20μg/ml) and hygromycin (50μg/ml) in culture treated 6-well plates (VWR) and cultured overnight at 23°C. Cells were infected (MOI 5) with mCerulean-producing *L. pneumophila* JR32 (pNP99), centrifuged (450 × *g*, 10 min, RT) and incubated at 25°C for 1 h. Subsequently, infected cells were washed three times with HL5 medium and incubated at 25°C for the time indicated. At given time points, infected cells (including supernatant) were collected from the 6-wells plate, centrifuged (500× *g*, 5 min, RT) and fixed with 4% PFA (Electron Microscopy Sciences) for 30min at RT. Following fixation, the cells were washed twice with PBS, transferred to a 16-wells μ-slide dish (Ibidi) and immobilized by adding a layer of PBS/0.5% agarose.

Image acquisition was performed using the confocal microscope Leica TCS SP8 X CLSM (HC PL APO CS2, objective 63×/1.4–0.60 oil; Leica Microsystems) with a scanning speed of 200Hz, bi-directional laser scan. Pictures were acquired with a pixel/voxel size close to the instrument’s Nyquist criterion of 43[×43×130nm (xyz). Images were deconvolved with Huygens professional version 19.10 software (Scientific Volume Imaging, http://svi.nl) using the CMLE algorithm, set to 10-20 iterations and quality threshold of 0.05.

### Imaging flow cytometry of infected *D. discoideum*

Processing of infected *D. discoideum* for IFC was performed as described [84, 85]. *D. discoideum* cells harboring the plasmids indicated were seeded in a 12-well plate containing HL5 medium, geneticin and hygromycin, and infected (MOI 5) with mPlum-producing *L. pneumophila* JR32, Δ*lepB* or Δ*sidC* (pAW014). The infection was synchronized by centrifugation (450 × *g*, 10 min, RT). The cells were incubated for 1 h at 25°C, washed three times with HL5 and incubated at 25°C. At the indicates time points cells were collected, centrifuged (450 × *g*, 5 min, RT) and fixed in 2 % PFA for 90 min on ice. Fixed infected amoebae were washed twice in phosphate-buffered saline (PBS) and resuspended in 20 µl ice cold PBS prior to IFC analysis.

At least 5000 cells were acquired using an imaging flow cytometer (ImageStramX MkII; Amnis) and analyzed with the IDEAS (v6.2) software (Amnis). Infected amoeba containing one intracellular *L. pneumophila* bacterium were gated and assessed for colocalization of GFP and mCherry (host) with mPlum produced by the bacteria. The software computes the IFC colocalization score (bright detail similarity), which is the log-transformed Pearson’s correlation coefficient of the localized bright spots with a radius of 3 pixels or less in two images and is used to quantify relative enrichment of a marker on the LCV. Data analysis was performed using GraphPad Prism.

### Comparative proteomics of purified LCVs

LCVs from *D. discoideum* amoebae were purified basically as described [86]. Briefly, *D. discoideum* Ax3 or Δ*sey1* producing CnxA-GFP (pAW016) was seeded in T75 flasks (3 per sample) one day before the experiment to reach 80% confluency. The amoebae were infected (MOI 50, 1 h) with *L. pneumophila* JR32 producing mCherry (pNP102) grown to stationary phase (21 h liquid culture). Subsequently, the cells were washed with SorC buffer and scraped in homogenization buffer (20 mM HEPES, 250 mM sucrose, 0.5 mM EGTA, pH 7.2) [45]. Cells were homogenized using a ball homogenizer (Isobiotec) with an exclusion size of 8 µm and incubated with an anti-SidC antibody followed by a secondary anti-rabbit antibody coupled to magnetic beads. The LCVs were separated in a magnetic field and further purified by a density gradient centrifugation step as described [87]. Three independent biological samples were prepared each for LCVs purified from *L. pneumophila*-infected *D. discoideum* Ax3 or Δ*sey1*.

LCVs purified by immuno-magnetic separation and density gradient centrifugation (fraction 4) were resolved by 1D-SDS-PAGE, the gel lanes were excised in ten equidistant pieces and subjected to trypsin digestion [88]. For the subsequent LC-MS/MS measurements, the digests were separated by reversed phase column chromatography using an EASY nLC 1000 (Thermo Fisher Scientific) with self-packed columns (OD 360 μm, ID 100 μm, length 20 cm) filled with 3 µm diameter C18 particles (Dr. Maisch, Ammerbuch-Entringen, Germany) in a one-column setup. Following loading/ desalting in 0.1% acetic acid in water, the peptides were separated by applying a binary non-linear gradient from 5-53% acetonitrile in 0.1% acetic acid over 82 min. The LC was coupled online to a LTQ Orbitrap Elite mass spectrometer (Thermo Fisher, Bremen, Germany) with a spray voltage of 2.5 kV. After a survey scan in the Orbitrap (r = 60,000), MS/MS data were recorded for the twenty most intensive precursor ions in the linear ion trap. Singly charged ions were not considered for MS/MS analysis. The lock mass option was enabled throughout all analyzes.

After mass spectrometric measurement, database search against a database of *Dictyostelium discoideum* and *Legionella pneumophila* Philadelphia downloaded Uniprot on 14/10/2019 (25,478 and 3,024 entries, respectively) as well as label-free quantification (LFQ) was performed using MaxQuant (version 1.6.7.0) (Cox, J and Mann, M (2008). MaxQuant enables high peptide identification rates, individualized p.p.b.-range mass accuracies and proteome-wide protein quantification. Nature Biotechnology 26, 1367–72). Common laboratory contaminants and reversed sequences were included by MaxQuant. Search parameters were set as follows: trypsin/P specific digestion with up to two missed cleavages, methionine oxidation and N-terminal acetylation as variable modification, match between runs with default parameters enabled. The FDRs (false discovery rates) of protein and PSM (peptide spectrum match) levels were set to 0.01. Two identified unique peptides were required for protein identification. LFQ was performed using the following settings: LFQ minimum ratio count 2 considering only unique for quantification.

Results were filtered for proteins quantified in at least two out of three biological replicates before statistical analysis. Here, two conditions were compared by a student’s t-test applying a threshold p-value of 0.01, which was based on all possible permutations. Proteins were considered to be differentially abundant if the log2-fold change was greater than |0.8|. “ON/OFF proteins” were defined as being identified in all bioreplicates of one strain whereas the protein was not identified in any replicate of the other strain.

### Statistical methods

Microscopy data analysis was performed using GraphPad Prism. The two-sample Student’s t-test (Mann-Whitnex test, no assumption of Gaussian distributions) was used. Probability values of less than 0.05, 0.01, and 0.001 were used to show statistically significant differences and are represented with *, **, or ***, respectively. The value of “n” represents the number of independent experiments performed or the number of analyzed cells per conditions (**Fig. 4B, Fig. 6B**). For the comparative proteomics, the summarized protein expression values were used for statistical testing of between condition differentially abundant proteins. Empirical Bayes moderated t-tests were applied, as implemented in the R/Bioconductor limma package.

## Data Availability

The MS proteomics data discussed in this publication have been deposited to the ProteomeXchange Consortium via the PRIDE [89] partner repository with the dataset identifier PXD034490 (Reviewer account details: Username; Password, JrGWYYBQ).

## Abbreviations

ATL: Atlastin

FFAT motif: two phenylalanines (FF) in an acidic tract motif

Icm/Dot: intracellular multiplication/defective organelle trafficking

IFC: imaging flow cytometry

LCV: *Legionella*-containing vacuole

GFP: green fluorescent protein

MCS: membrane contact sites

OSBP: oxysterol binding protein

T4SS: type IV secretion system

VAMP: vesicle-associated membrane protein

Vap: VAMP-associated protein.

## Acknowledgements

We would like to thank Markus Maniak for providing the deletion construct for *osbH*, Leoni Swart, Xiaoli Ma, Deise Schäfer, and Iris Hube for help with cloning, and Sebastian Grund for technical support with the preparation of MS samples. Work in the group of H.H. was supported by the Swiss National Science Foundation (SNF; 31003A_175557, 310030_207826). Work in the group of D.B. was supported by the Federal Ministry of Education and Research (BMBF; grant 031A410B). Work in the group of C.B. was supported by the DFG and the SFB944 (grant SFB 944/3-P25). Work in the group of F.L. was supported by the Région Occitanie, the Centre National de la Recherche Scientifique (CNRS) and the University of Montpellier (UM). The authors declare no conflict of interest.

## Supplementary Figures

**Figure S1.** Phylogenetic tree of short OSBPs from *D. discoideum*, *Saccharomyces cerevisiae* and humans. Sequences of all proteins were either derived from dictybase.org or uniprot and aligned with MAFFT (https://mafft.cbrc.jp) using the G-INS-I strategy, unalignlevel 0.8 and “leave gappy regions” to generate a phylogenetic tree in phylo.ilo using NJ conserved sites and the JTT substitution model. Numbers on the branches indicate bootstrap support for nodes from 100 bootstrap replicates.

**Figure S2.** *Dictyostelium discoideum* MCS components localize to LCVs and/or ER. (**A**) Dually labeled *D. discoideum* amoebae producing MCS components fused to GFP and either CnxA-mCherry (pAW016), P4C-mCherry (pWS032), or AmtA-mCherry were infected (MOI 5, 2 h) with mCerulean-producing *L. pneumophila* JR32 (pNP99), fixed with 4 % PFA, and imaged by confocal fluorescence microscopy. Single channels are shown. scale bars: 3 μm. The Pearson’s correlation coefficient of uninfected cells was generated using Coloc 2 from Fiji (ImageJ) and is shown for MCS components fused to GFP with respect to (**B**) CnxA-mCherry, (**C**) P4C-mCherry, or (**D**) AmtA-mCherry. Data represent mean and SEM of three independent experiments (*P<0.05; ***P<0.001).

**Figure S3.** Construction of *D. discoideum* deletion mutants. Schematic representation of the strategies followed for the construction of *D. discoideum* strains lacking (**A**) *osbG*, (**B**) *osbH*, (**C**) *osbK*, or (**D**) *vap*, and PCR-based validation.

**Figure S4.** MCS components modulate replication of *L. pneumophila* in *D. discoideum* and mammalian cells. (**A**) *D. discoideum* Ax3, or Δ*vap*, Δ*osbG,* Δ*osbH*, or Δ*osbK* mutants producing GFP-Sac1 (pLS037) or GFP-Sac1_ΔTMD (pSV034) were infected (MOI 1) with mCherry-producing *L. pneumophila* JR32 (pNP102), and intracellular replication was assessed by colony-forming units (CFU). Mean and SEM of three independent experiments are shown (*P<0.05; **P<0.01; ***P<0.001). To improve clarity, the different mutant strains are shown in separate graphs, each depicting the same data for Ax3/GFP-Sac1. (**B**) Cytotoxicity toward A549 cells of oligonucleotides targeting Arf1 and Sac1 (10 nM siRNA, 48 h) was determined by the Zombie Aqua fixable viability kit (BioLegend) using flow cytometry. Percentage of Zombie-positive cells is shown (means and SEM of triplicate experiments). Untreated cells were used as a negative control, and treatment with 70% EtOH for 1 h served as positive control for cell death. (**C**) A549 epithelial cells were treated with oligonucleotides targeting Sac1 (10 nM siRNA, 48 h), and the efficiency of protein depletion was assessed by Western blot (WB) with the antibodies indicated. Qiagen AllStars unspecific oligonucleotides (“Scrambled”, “Scr”) were used to control for off-target effects, and GAPDH served as WB loading control. Data are representative of two independent experiments.

**Figure S5.** Vap, OSBP11, and Sac1 promote expansion of PtdIns(4)*P*-positive LCVs. (**A**) Dually labeled *D. discoideum* Ax3, Δ*vap*, Δ*osbG,* Δ*osbH*, Δ*osbK* or Δ*sey1* mutants producing P4C-mCherry (pWS032) and CnxA-GFP (pAW016), or Ax3 producing P4C-mCherry and either GFP-Sac1 (pLS037), GFP-Sac1_C383S (pSV015) or GFP-Sac1_ΔTMD (pSV034) were infected (MOI 5, 2-16 h) with mCerulean-producing *L. pneumophila* JR32 (pNP99) and fixed with 4 % PFA. Single channels for the analyzed time points are shown. Scale bars: 3 μm. (**B**) LCV area was measured using ImageJ (n=100-200 per condition from 3 independent biological replicates). Means and SEM of single cells are shown (*P<0.05; **P<0.01; ***P<0.001). The data for Ax3/GFP-Sac1 and Ax3/GFP Sac1_ΔTMD is also shown in Fig. 4B.

**Figure S6.** Localization of GFP-Sac1 in *D. discoideum* Ax3 or strains lacking MCS components. Dually labeled *D. discoideum* Ax3, Δ*vap*, Δ*osbG,* Δ*osbH*, Δ*osbK* or Δ*sey1* mutants producing P4C-mCherry (pWS032) and GFP-Sac1 (pLS037) were infected (MOI 5, 2-16 h) with mCerulean-producing *L. pneumophila* JR32 (pNP99) and fixed with 4 % PFA. Single channels for the analyzed time points are shown. Scale bars: 3 μm.

**Figure S7.** Localization of GFP-MCS components in *D. discoideum* wild-type producing Sac1 or Sac1_ΔTMD. Dually labeled *D. discoideum* Ax3 producing GFP fusions of MCS proteins and either mCherry-Sac1 (pSV044) or mCherry-Sac1_ΔTMD (pSV045) were infected (MOI 5, 2 h) with mCerulean-producing *L. pneumophila* JR32 (pNP99) and fixed with 4 % PFA. Single channels for the analyzed time points are shown. Scale bars: 3 μm.

**Figure S8.** The *L. pneumophila* effectors LepB and SidC determine PtdIns(4)*P* decoration and expansion of LCVs. Dually labeled *D. discoideum* Ax3 producing P4C-mCherry (pWS032) and either CnxA-GFP (pAW016), GFP-Sac1 (pLS037), or GFP-Sac1_ΔTMD (pSV034) were infected (MOI 5, 1-8 h) with mCerulean-producing *L. pneumophila* JR32, Δ*lepB* or Δ*sidC* (pNP99) and fixed with 4 % PFA. Single channels for the analyzed time points are shown. Scale bars: 3 μm.

**Table S1.** Comparative proteomics of LCVs from *D. discoideum* Ax3 and Δ*sey1*.

**Table S2.** Cells, bacterial strains, and plasmids used in this study.

**Table S3.** Oligonucleotides used in this study.

## Notes

### Competing Interest Statement

The authors have declared no competing interest.

### Summary of Updates

Supplemental Information has been uploaded.

