## Supplementary Information for "The *Legionella*-driven PtdIns(4)*P* gradient at LCV-ER membrane contact sites promotes Vap-, OSBP- and Sac1-dependent pathogen vacuole remodeling"

**Running title:** LCV remodeling through ER membrane contact sites

**Key words:** Amoeba, atlastin, *Dictyostelium discoideum*, large fusion GTPase, host-pathogen interaction, *Legionella pneumophila*, Legionnaires' disease, oxysterol binding protein, pathogen vacuole.

**Abbreviations:** ATL, Atlastin, FFAT motif, two phenylalanines (FF) in an acidic tract motif; Icm/Dot, intracellular multiplication/defective organelle trafficking; IFC, imaging flow cytometry; LCV, *Legionella*-containing vacuole; GFP, green fluorescent protein; MCS, membrane contact sites; OSBP, oxysterol binding protein; T4SS, type IV secretion system; VAMP, vesicle-associated membrane protein; Vap, VAMP-associated protein.

### Supplementary Figures

**Figure S1**

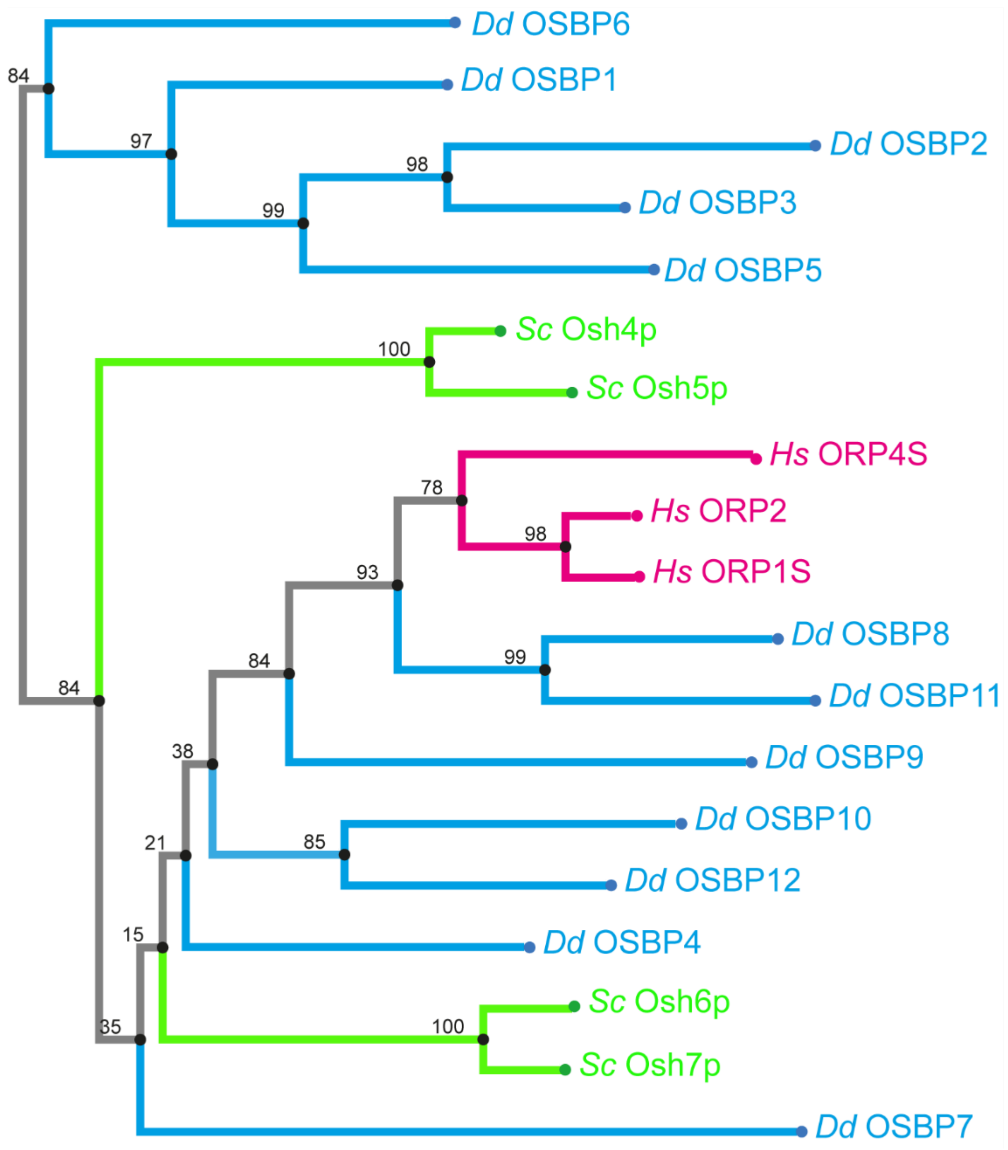


**Fig. S1.** Phylogenetic tree of short OSBPs from *D. discoideum*, *Saccharomyces cerevisiae* and humans.

Sequences of all proteins were either derived from dictybase.org or uniprot and aligned with MAFFT (https://mafft.cbrc.jp) using the G-INS-I strategy, unalignlevel 0.8 and “leave gappy regions” to generate a phylogenetic tree in phylo.ilo using NJ conserved sites and the JTT substitution model. Numbers on the branches indicate bootstrap support for nodes from 100 bootstrap replicates.

#### Figure S2

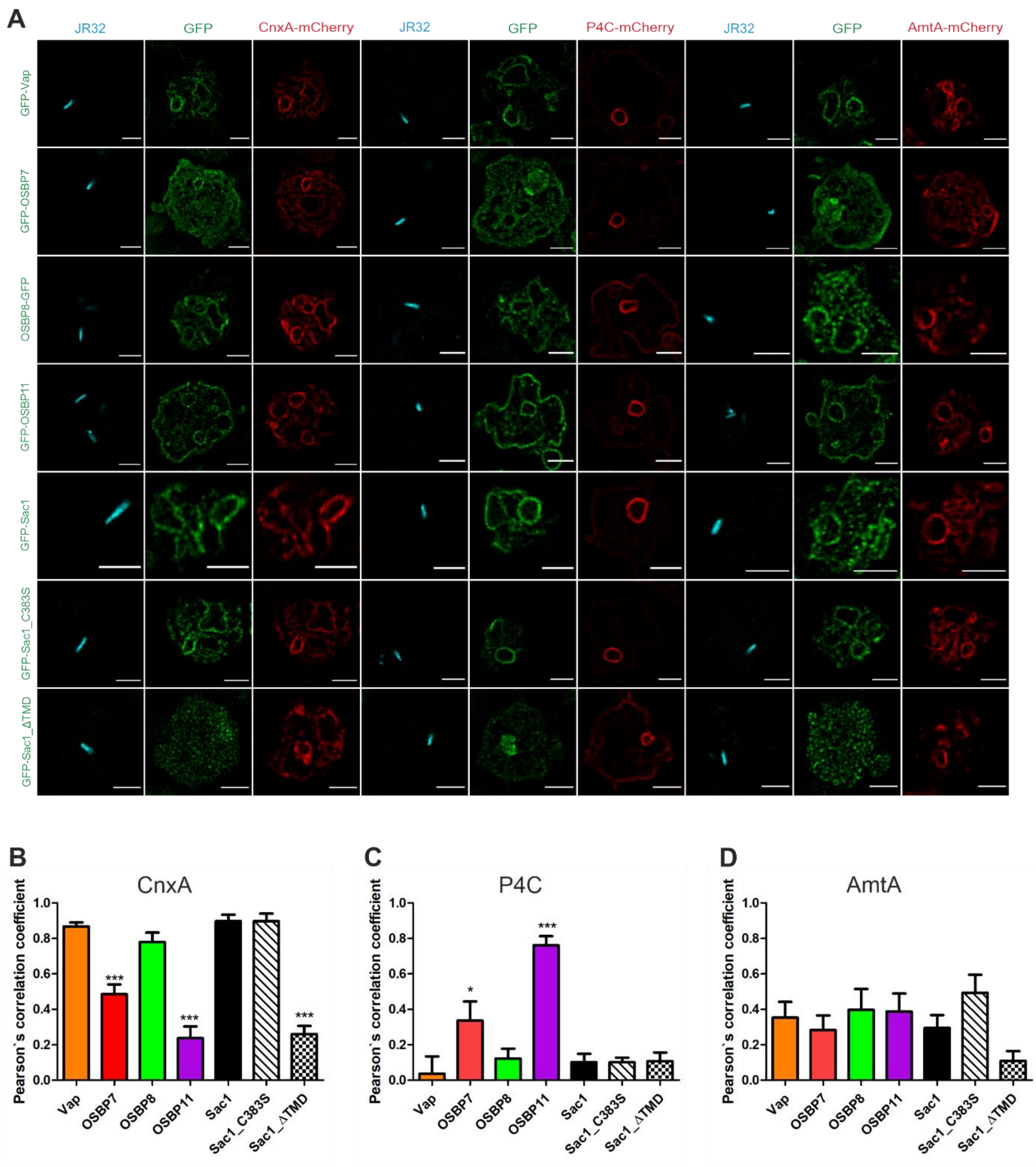


**Fig. S2.** *Dictyostelium discoideum* MCS components localize to LCVs and/or ER.
(**A**) Dually labeled *D. discoideum* amoebae producing MCS components fused to GFP and either CnxA-mCherry (pAW016), P4C-mCherry (pWS032), or AmtA-mCherry were infected (MOI 5, 2 h) with mCerulean-producing *L. pneumophila* JR32 (pNP99), fixed with 4 % PFA, and imaged by confocal fluorescence microscopy. Single channels are shown. scale bars: 3 µm. The Pearson's correlation coefficient of uninfected cells was generated using Coloc 2 from Fiji (ImageJ) and is shown for MCS components fused to GFP with respect to (**B**) CnxA-mCherry, (**C**) P4C-mCherry, or (**D**) AmtA-mCherry. Data represent mean and SEM of three independent experiments (*P<0.05; ***P<0.001).

**Figure S3**

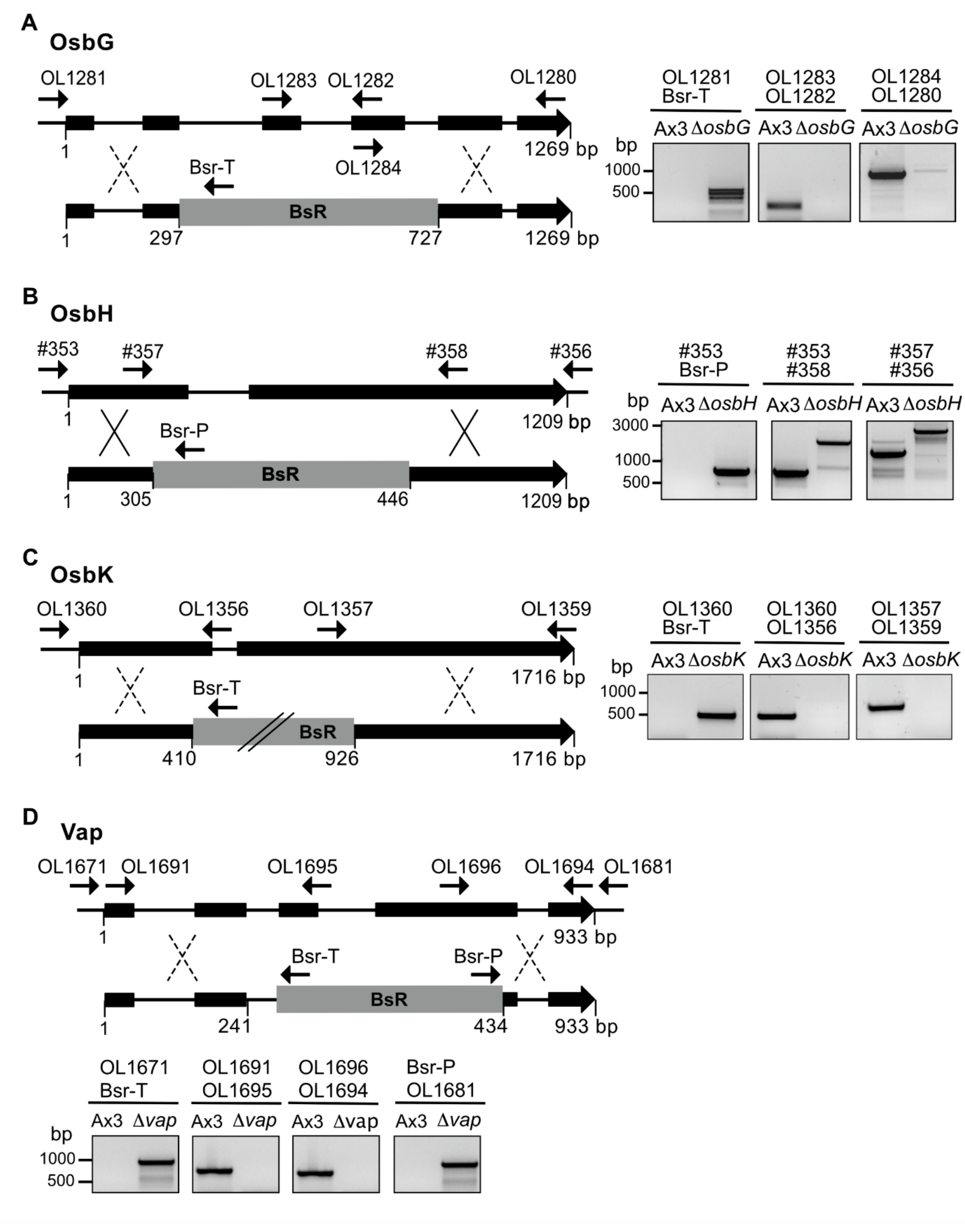


**Fig. S3.** Construction of *D. discoideum* deletion mutants. Schematic representation of the strategies followed for the construction of *D. discoideum* strains lacking (**A**) *osbG*, (**B**) *osbH*, (**C**) *osbK*, or (**D**) *vap*, and PCR-based validation.

**Figure S4**

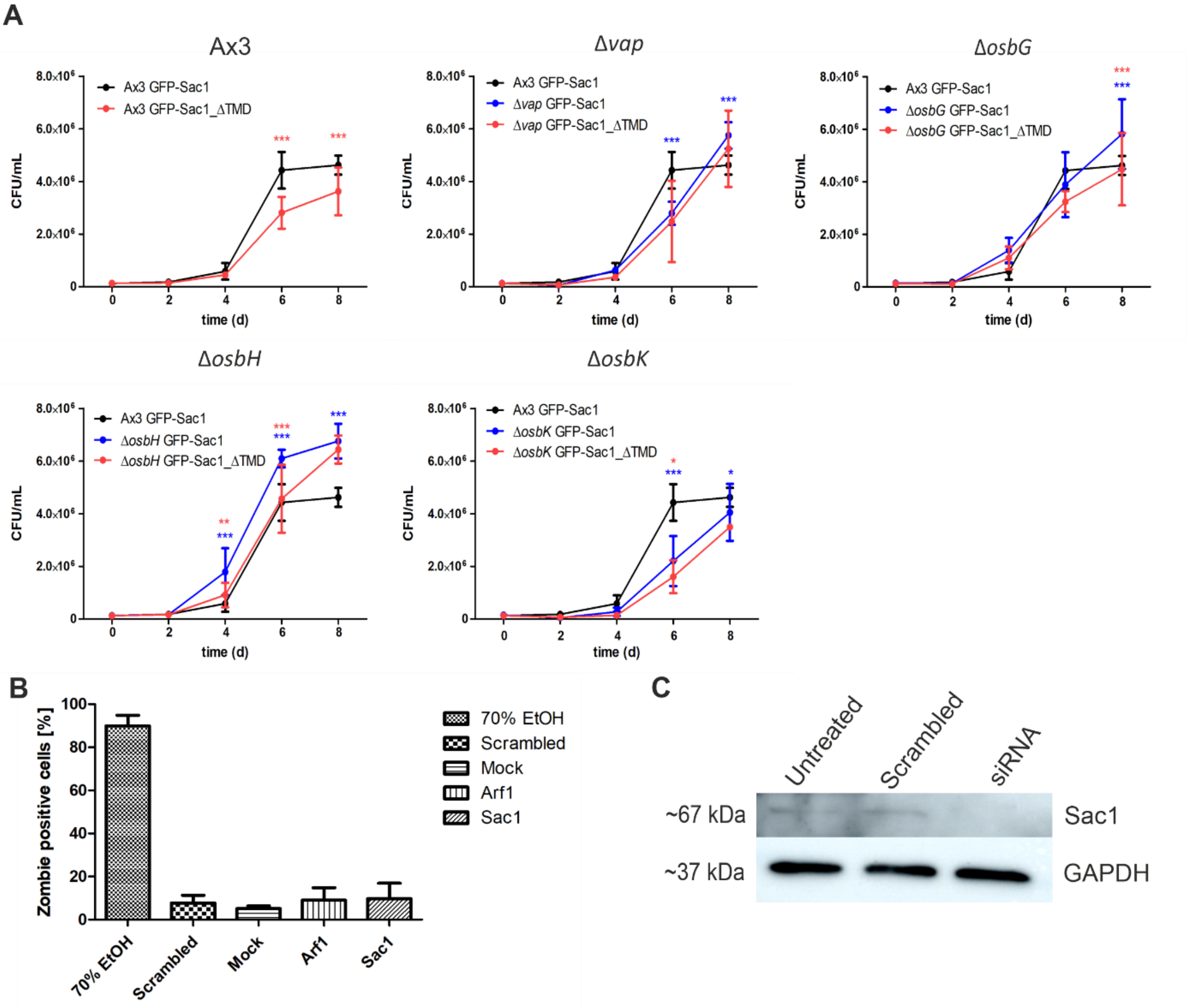


**Fig. S4.** MCS components modulate replication of *L. pneumophila* in *D. discoideum* and mammalian cells.

(**A**) *D. discoideum* Ax3, or Δ*vap*, Δ*osbG*, Δ*osbH*, or Δ*osbK* mutants producing GFP-Sac1 (pLS037) or GFP-Sac1_ΔTMD (pSV034) were infected (MOI 1) with mCherry-producing *L. pneumophila* JR32 (pNP102), and intracellular replication was assessed by colony-forming units (CFU). Mean and SEM of three independent experiments are shown (*P<0.05; **P<0.01; ***P<0.001). To improve clarity, the different mutant strains are shown in separate graphs, each depicting the same data for Ax3/GFP-Sac1. (**B**) Cytotoxicity toward A549 cells of oligonucleotides targeting Arf1 and Sac1 (10 nM siRNA, 48 h) was determined by the Zombie Aqua fixable viability kit (BioLegend) using flow cytometry. Percentage of Zombie-positive cells is shown (means and SEM of triplicate experiments). Untreated cells were used as a negative control, and treatment with 70% EtOH for 1 h served as positive control for cell death. (**C**) A549 epithelial cells were treated with oligonucleotides targeting Sac1 (10 nM siRNA, 48 h), and the efficiency of protein depletion was assessed by Western blot (WB) with the antibodies indicated. Qiagen AllStars unspecific oligonucleotides ("Scrambled", "Scr") were used to control for off-target effects, and GAPDH served as WB loading control. Data are representative of two independent experiments.

#### Figure S5

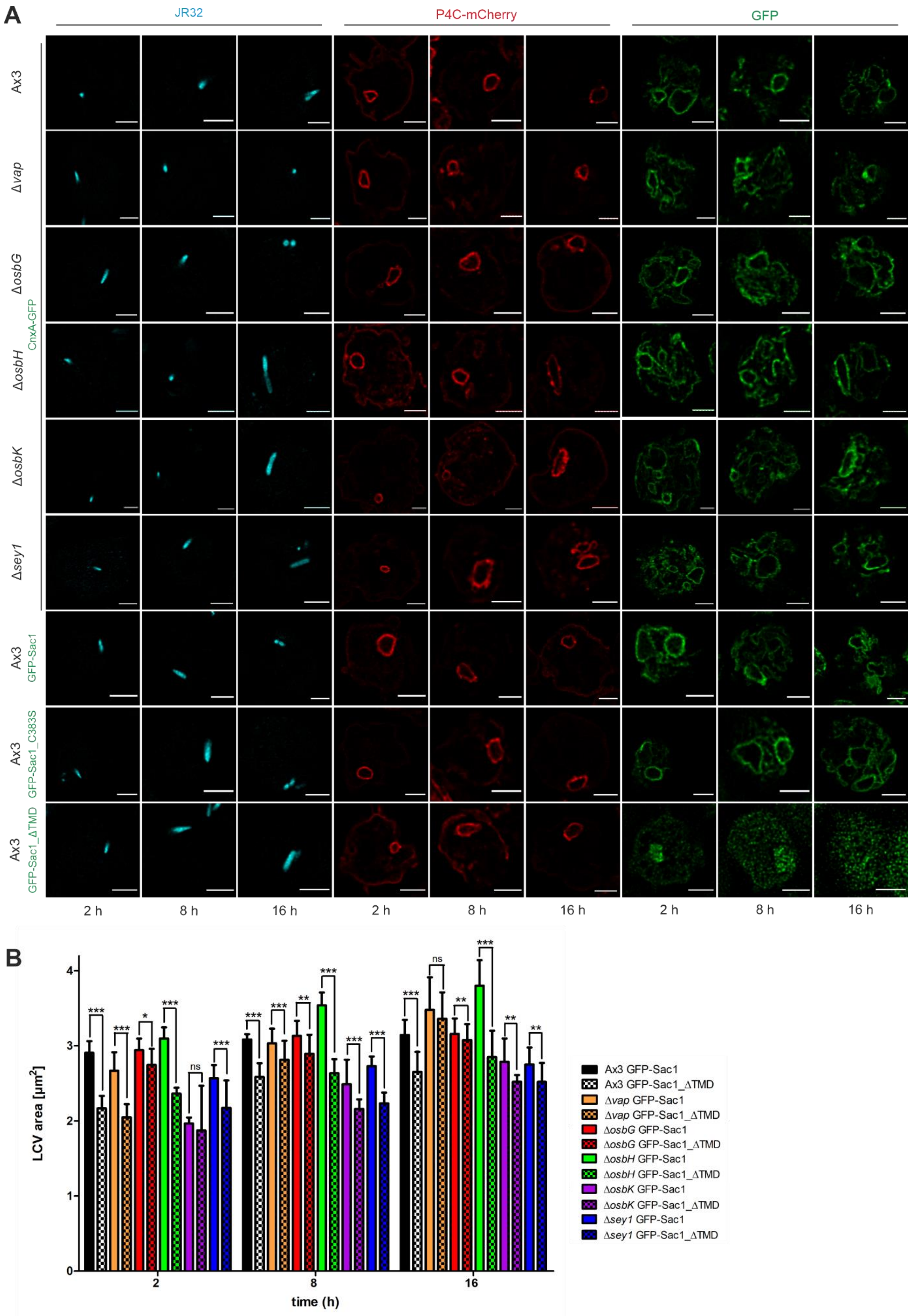

A
JR32
P4C-mCherry
GFP
Ax3
Δvap
ΔosbG
ΔosbH
ΔosbK
Δsey1
CnxA-GFP
Ax3 GFP-Sac1
Ax3 GFP-Sac1_C383S
Ax3 GFP-Sac1_ΔTMD
2 h
8 h
16 h
B
LCV area [µm²]
time (h)
2
8
16
ns
Ax3 GFP-Sac1
Ax3 GFP-Sac1_ΔTMD
Δvap GFP-Sac1
Δvap GFP-Sac1_ΔTMD
ΔosbG GFP-Sac1
ΔosbG GFP-Sac1_ΔTMD
ΔosbH GFP-Sac1
ΔosbH GFP-Sac1_ΔTMD
ΔosbK GFP-Sac1
ΔosbK GFP-Sac1_ΔTMD
Δsey1 GFP-Sac1
Δsey1 GFP-Sac1_ΔTMD

**Fig. S5** (overleaf)**.** Vap, OSBP11, and Sac1 promote expansion of PtdIns(4)*P*-positive LCVs. (**A**) Dually labeled *D. discoideum* Ax3, Δ*vap*, Δ*osbG*, Δ*osbH*, Δ*osbK* or Δ*sey1* mutants producing P4C-mCherry (pWS032) and CnxA-GFP (pAW016), or Ax3 producing P4C-mCherry and either GFP-Sac1 (pLS037), GFP-Sac1_C383S (pSV015) or GFP-Sac1_ΔTMD (pSV034) were infected (MOI 5, 2-16 h) with mCerulean-producing *L. pneumophila* JR32 (pNP99) and fixed with 4 % PFA. Single channels for the analyzed time points are shown. Scale bars: 3 μm. (**B**) LCV area was measured using ImageJ (n=100-200 per condition from 3 independent biological replicates). Means and SEM of single cells are shown (*P<0.05; **P<0.01; ***P<0.001). The data for Ax3/GFP-Sac1 and Ax3/GFP Sac1_ΔTMD is also shown in Fig. 4B.

**Fig. S6** (overleaf)**.** Localization of GFP-Sac1 in *D. discoideum* Ax3 or strains lacking MCS components.

Dually labeled *D. discoideum* Ax3, Δ*vap*, Δ*osbG*, Δ*osbH*, Δ*osbK* or Δ*sey1* mutants producing P4C-mCherry (pWS032) and GFP-Sac1 (pLS037) were infected (MOI 5, 2-16 h) with mCerulean-producing *L. pneumophila* JR32 (pNP99) and fixed with 4 % PFA. Single channels for the analyzed time points are shown. Scale bars: 3 μm.

**Figure S6**

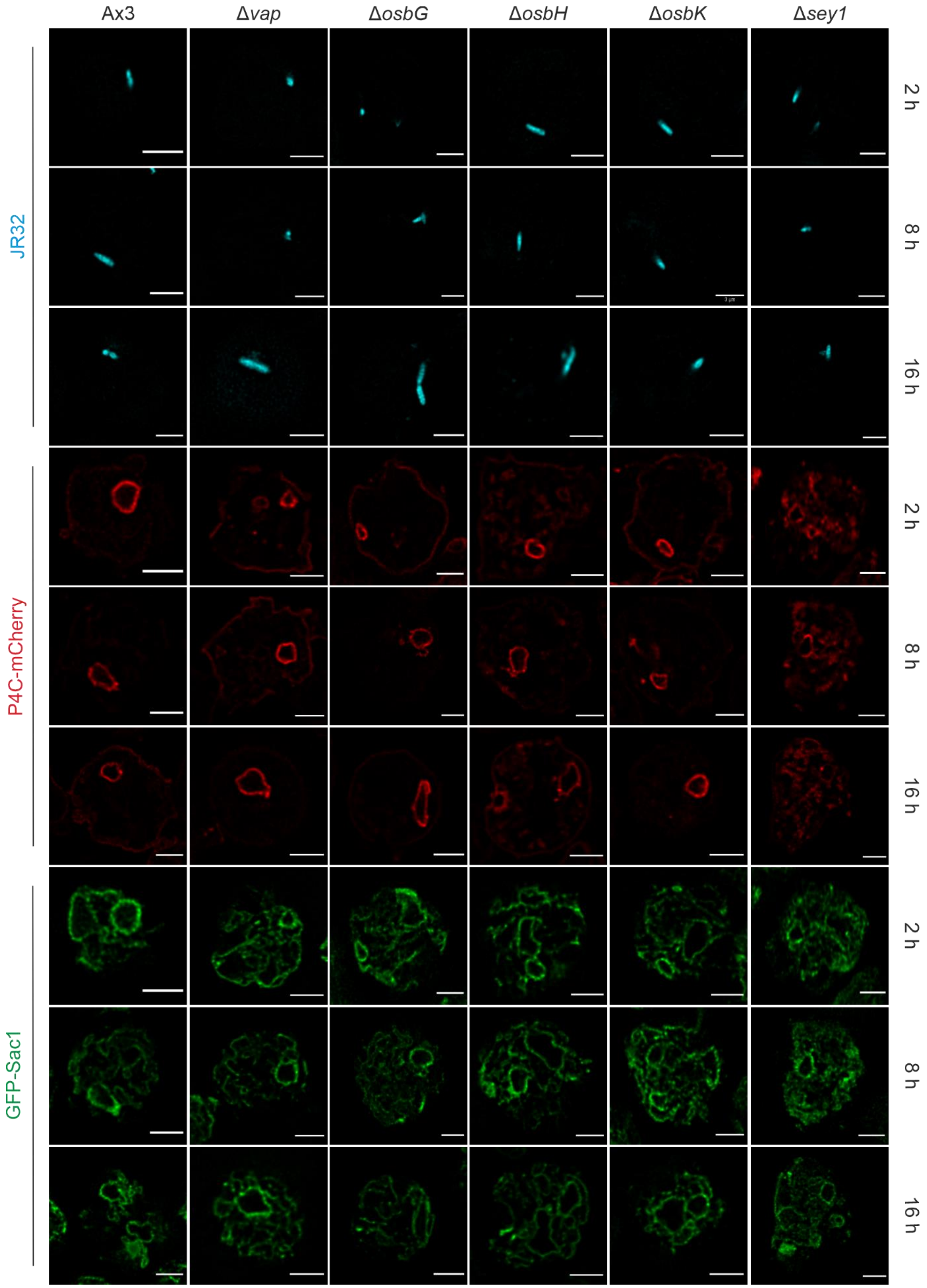

Ax3
Δvap
ΔosbG
ΔosbH
ΔosbK
Δsey1
JR32
P4C-mCherry
GFP-Sac1
2 h
8 h
16 h
2 h
8 h
16 h
2 h
8 h
16 h

**Figure S7**

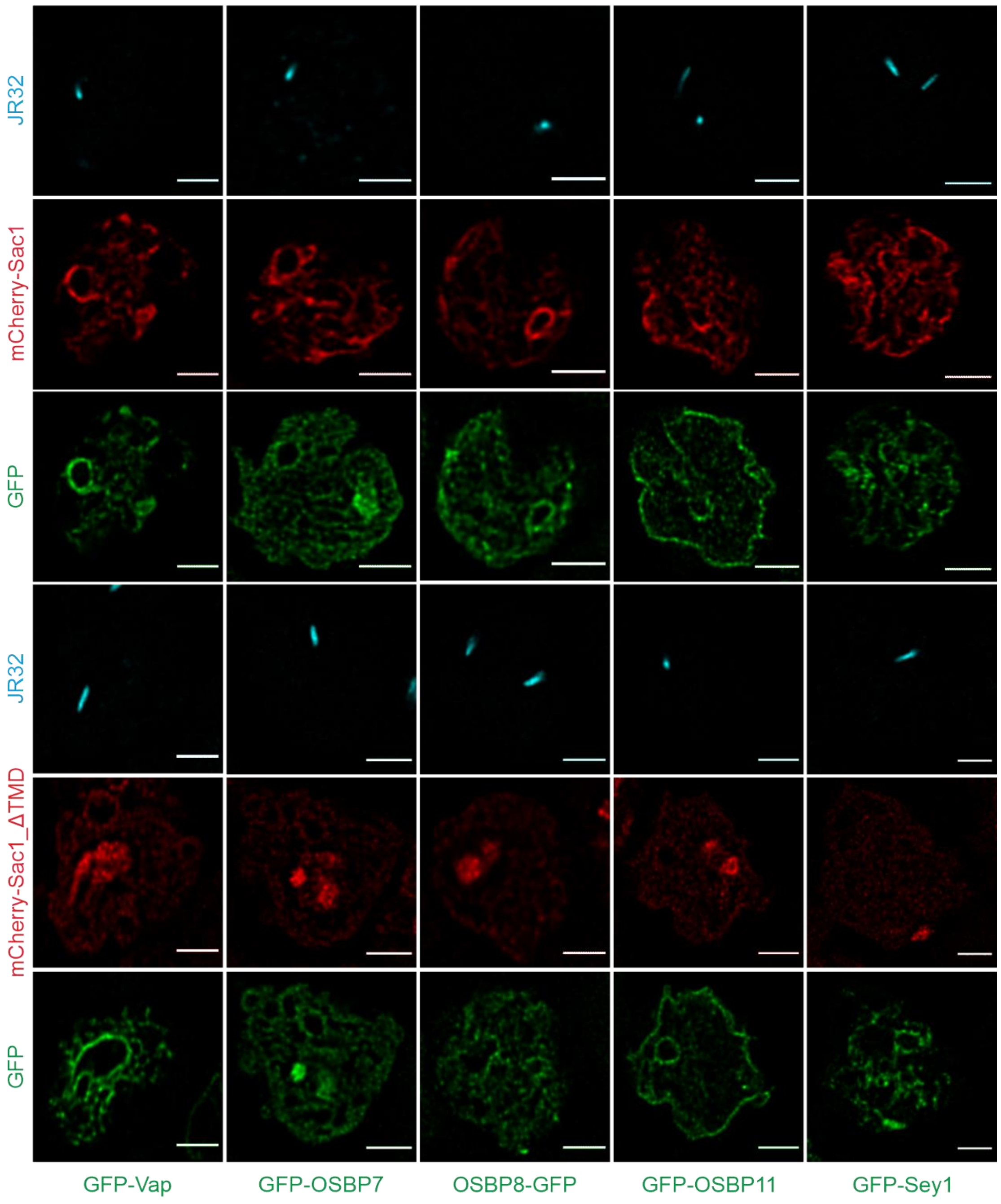


**Fig. S7.** Localization of GFP-MCS components in *D. discoideum* wild-type producing Sac1 or Sac1_ΔTMD.

Dually labeled *D. discoideum* Ax3 producing GFP fusions of MCS proteins and either mCherry-Sac1 (pSV044) or mCherry-Sac1_ΔTMD (pSV045) were infected (MOI 5, 2 h) with mCerulean-producing *L. pneumophila* JR32 (pNP99) and fixed with 4 % PFA. Single channels for the analyzed time points are shown. Scale bars: 3 µm.

**Figure S8**

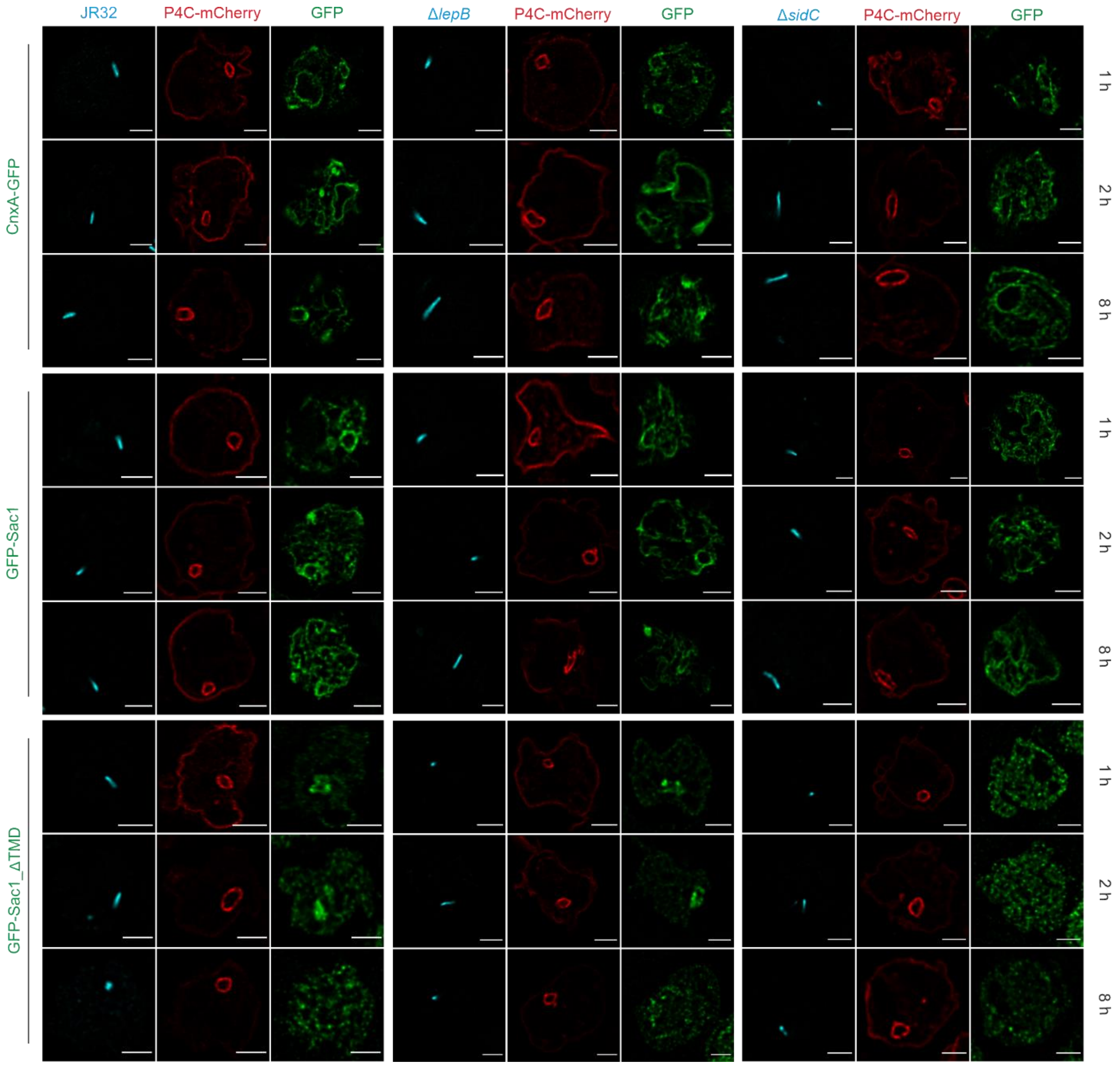


**Fig. S8.** The *L. pneumophila* effectors LepB and SidC determine PtdIns(4)*P* decoration and expansion of LCVs.

Dually labeled *D. discoideum* Ax3 producing P4C-mCherry (pWS032) and either CnxA-GFP (pAW016), GFP-Sac1 (pLS037), or GFP-Sac1_ΔTMD (pSV034) were infected (MOI 5, 1-8 h) with mCerulean-producing *L. pneumophila* JR32, Δ*lepB* or Δ*sidC* (pNP99) and fixed with 4 % PFA. Single channels for the analyzed time points are shown. Scale bars: 3 µm.

### Supplementary Tables

**Table S2.** Cells, bacterial strains, and plasmids used in this study.

| Strain or plasmid | Relevant properties [a] | Reference |
|---|---|---|
| ***D. discoideum*** | | |
| Ax3 | Parental strain | [1] |
| Δ*osbG* | Ax3, insertion in gene DDB_G0283035, $Bls^R$ | This study |
| Δ*osbH* | Ax3, insertion in gene DDB_G0283709, $Bls^R$ | This study |
| Δ*osbK* | Ax3, insertion in gene DDB_G0288817, $Bls^R$ | This study |
| Δ*sey1* | Ax3, insertion in gene DDB_G0279823, $Bls^R$ | [2] |
| Δ*vap* | Ax3, insertion in gene DDB_G0278773, $Bls^R$ | This study |
| **Mammalian cells** | | |
| A549 | Human adenocarcinoma alveolar epithelial cells | ATCC CCL-185 |
| RAW 264.7 | Murine macrophages | ATCC TIB-71 |
| ***E. coli*** | | |
| TOP10 | | Invitrogen |
| ***L. pneumophila*** | | |
| CR001 (Δ*sidC*) | JR32 *sidC-sdcA::*$Kan^R$ *(ΔsidC-sdcA)* | [3] |
| GS3011 (Δ*icmT*) | JR32 *icmT3011*:: $Kan^R$ | [4] |
| IH03 (Δ*lepB*) | JR32 *lepB::*$Kan^R$ *(ΔlepB)* | This study |
| JR32 | *L. pneumophila* Philadelphia-1, serogroup 1, salt-sensitive isolate of AM511 | [5] |
| **Plasmids** | | |
| pAW012 | pDM1044, calnexin-mCherry, $Hyg^R$, $Amp^R$ | [6] |
| pAW014 | pMMB207C, Δ$lacI^q$, mPlum (constitutive), $Cam^R$ | [6] |
| pAW016 | pDM323, calnexin-GFP, $G418^R$, $Amp^R$ | [7] |
| pDM317 | *Dictyostelium* expression vector, extra-chromosomal, N-terminal GFP, $G418^R$, $Amp^R$ | [8] |
| pDM323 | *Dictyostelium* expression vector, extra-chromosomal, C-terminal GFP, $G418^R$, $Amp^R$ | [8] |
| pDM1042 | *Dictyostelium* expression vector, extra-chromosomal, N-terminal mCherry, $Hyg^R$, $Amp^R$ | [9] |
| pDM1044 | *Dictyostelium* expression vector, extra-chromosomal, C-terminal mCherry, $Hyg^R$, $Amp^R$ | [9] |
| pDM1044-AmtA-mCherry | pDM1044, AmtA-mCherry, $Hyp^R$, $Amp^R$ | [9] |

| | | |
|---|---|---|
| pFL1517 | pBluescript, Δ*osbG*, $Bls^R$, $Amp^R$ | This work |
| pFL1542 | pBluescript, Δ*osbK*, $Bls^R$, $Amp^R$ | This work |
| pFL1545 | pBluescript, Δ*vap*, $Bls^R$, $Amp^R$ | This work |
| pMIB39 | pDM317, GFP-OSBP11 (*osbK*), $G418^R$, $Amp^R$ | This work |
| pMIB41 | pDM317, GFP-Vap, $G418^R$, $Amp^R$ | This work |
| pMIB87 | pDM317, GFP-OSBP7 (*osbG*), $G418^R$, $Amp^R$ | This work |
| pMIB89 | pDM323, OSBP8-GFP (*osbH*), $G418^R$, $Amp^R$ | This work |
| pMM629 | pGEM-T Easy, Δ*osbH*, $Bls^R$, $Amp^R$ | Gift from M. Maniak |
| pLAW344 | *oriT* (RK2), *oriR* (ColE1), *sacB*, $Cam^R$, $Amp^R$ | [10] |
| pLS037 | pDM317, GFP-Sac1, $G418^R$, $Amp^R$ | This work |
| pIH29 | pLAW344-upstream *lepB*-$Kan^R$-downstream *lepB* | This work |
| pNP99 | pMMB207C, Δ*lacI*$^q$, mCerulean (constitutive), $Cam^R$ | [6] |
| pNP102 | pMMB207C, Δ*lacI*$^q$, mCherry (constitutive), $Cam^R$ | [6] |
| pNT28 | pMMB207C, Δ*lacI*$^q$, GFP (constitutive), $Cam^R$ | [11] |
| pSV015 | pDM317, GFP-Sac1_C383S, $G418^R$, $Amp^R$ | This work |
| pSV034 | pDM317, GFP-Sac1_1-517 (Sac1_ΔTMD), $G418^R$, $Amp^R$ | This work |
| pSV044 | pDM1042, mCherry-Sac1, $Hyg^R$, $Amp^R$ | This work |
| pSV045 | pDM1042 mCherry-Sac1_1-517 (Sac1_ΔTMD), $Hyg^R$, $Amp^R$ | This work |
| pWS032 | pDM1044, P4C-mCherry, $Hyg^R$, $Amp^R$ | [6] |

[a] Abbreviations: Amp, ampicillin; Bls, blasticidin S; Cam, chloramphenicol; Hyg, hygromycin; Kan, kanamycin; G418, geneticin.

**Table S3.** Oligonucleotides used in this study.

**Cloning**

| **Oligo** | **Sequence (5' - 3')** [a] | **Comments** |
|---|---|---|
| Bsr-T | TCAAAAAGATAAAGCTGACCCGAAAGC | 3' *Bls* (fo) |
| Bsr-P | CGCTACTTCTACTAATTCTAGA | 5' *Bls* (re) |
| oIH011 | AAAAACGCTCTAGATCTCAGGAACCATCAACACG | 5'flanking seq of *lepB* (fo), XbaI |
| oIH012 | AAAAACGCTCTAGACACCAGTTCACTCCATAAGG | 3'flanking seq of *lepB* (fo), XbaI |
| oIH013 | AAAAACGCGGATCCAATGACTCTGACCTAAAACAAC | 5'flanking seq of *lepB* (re), BamHI |
| oIH014 | AAAAACGCGGATCCTCTTCTCTTTTTGGCATAGG | 3'flanking seq of *lepB* (re), BamHI |
| oL1256 | AATAATTCTAGATTTATGTCAAATTTTTTCAAAAAGTTAGTTA AAAAG | 5' *osbK* (fo), XbaI |
| oL1277 | AATATAGGATCCTAAATGGAGGCCGATCCGAGCTTAGTTTCT | 5' *osbG* (fo), BamHI |
| oL1278 | AATATAAAGCTTTTAATTTCCATCTCTAACACAAGAAAAAAA GTATTGAATAATAATTGC | 5' *osbG* (re), HindIII |
| oL1279 | AATATAAAGCTTTAATATGTTAAAGGTGAAGTTTCAGTTTAT AATACAGAATTAGAAGTT | 3' *osbG* (fo), HindIII |
| oL1280 | AATATACTCGAGTTAATTACTACCACTTGCAGCATCAGAAGCA | 3' *osbG* (re), XhoI |
| oL1281 | AAATCCCAACACACTCGCGTGTAAATAAT | Δ*osbG* screening (fo) |
| oL1282 | AGTTCATATGGAATTTAGCTGAAAATTGTACATTACA | Δ*osbG* screening (re) |
| oL1283 | CACATCATCCACCATTAACAGCATTCAA | Δ*osbG* screening (fo) |
| oL1284 | TGTAATGTACAATTTTCAGCTAAATTCCATATGAACT | Δ*osbG* screening (fo) |
| oL1356 | ACATTAATCTATCGATTGAATCAGAACAAG | 5' *osbK* (re) |
| oL1357 | GTACAAGGTGAAATTCTTGATTCAAAGGG | Δ*osbK* screening (fo) |
| oL1358 | AATAATAAGCTTGCACGTGTACATATTAATGGTAAATGGGAT | 3' *osbK* (fo), HindIII |
| oL1359 | ATTATTCTCGAGTTATCTACCACTATGACCAATTGTAGGACT | 3' *osbK* (re), XhoI |
| oL1360 | TTTAATTAAATACTTTATAAGGTACCTCTGGTA | Δ*osbK* screening (fo) |
| oL1691 | AATATAGGATCCATGTCAAATAATGTTTCATCCAGTCCA | 5' *vap* (fo), BamHI |
| oL1692 | ATTATTAAGCTTCTAAATTTTGAAATTTGATTTATATGTTTAA AAATG | 5' *vap* (re), HindIII |
| oL1693 | ATTATTAAGCTTCATCAAATCCATCAAGTAACAGCAC | 3' *vap* (fo), HindIII |
| oL1694 | AATAATCTCGAGCATCAATTTATTAAATTAATTTACCAAAAAT AAAAG | 3' *vap* (re), XhoI |

| | | |
|---|---|---|
| oL1695 | CAATTGAAAGAATTTGAAATTTATCTTTGG | Δ*vap* screening (re) |
| oL1696 | GGACAAATTCACCACCAAATTTAATCA | Δ*vap* screening (fo) |
| oL1671 | AGCGCGTCTCCAATGGAATTCGTGGAAGGGAAAAC | Δ*vap* screening (fo) |
| oL1681 | ATTATTCTCGAGGTGTGGGGTGATAG | Δ*vap* screening (re) |
| oLS039 | AAAAGCGAGATCTAAAATGAATATTGAATTAATTAATAC | 5’ *sac1* (fo), BglII |
| oLS040 | TTTTTCGCACTAGTGTTTTTATAAATTGAATC | 3’ *sac1* (re), SpeI |
| oM353 | CAATACCAATAGATTTTATATCATTAC | Δ*osbH* screening (fo) |
| oM356 | CAGCGGAAATTGAATGAATAAATTG | Δ*osbH* screening (re) |
| oM357 | GCCTCAAAACAAGATAGC | Δ*osbH* screening (fo) |
| oM358 | CCTCTGATGAGTTACCATAG | Δ*osbH* screening (re) |
| oMIB38 | CGTCCGGAAAAATGTCAAATTTTTTCAAAAAGTTAG | 5’ *osbK* (fo), Kpn2I |
| oMIB39 | CCACTAGTTTATCTACCACTATGACCAATTG | 3’ *osbK* (re), BcuI |
| oMIB12 | CCACTAGTTTAAATTAATTTACCAAAAATAAAAG | 3’ *vap* (re), BcuI |
| oMIB14 | CGAGATCTAAAATGTCAAATAATGTTTCATCC | 5’ *vap* (fo), BglII |
| oMIB18 | CCACTAGTTTAATTACTACCACTTGCAGC | 3’ *osbG* (re), BcuI |
| oMIB20 | CGAGATCTAAAATGGAGGCCGATCCG | 5’ *osbG* (fo), BglII |
| oMIB21 | CGAGATCTAAAATGTTTTCAGGAGCATTG | 5’ *osbH* (fo), BglII |
| oMIB22 | CCACTAGTTTAATTTGAAGCTGCTGC | 3’ *osbH* (re), BcuI |
| oSV159 | GTTTTCCGTACGAATAGCATCGATAATTTAGAC | Sac1_C383S (fo) |
| oSV161 | ACCTTGTTGTTTTTGAACGATTTTACC | Sac1_C383S (re) |
| oSV227 | GAATTATATAAAGGTGGTTCAGGAGGTAGTAGATCTAAAATG AATATTGAATTAATTAATAC | Sac1_ΔTMD (fo) |
| oSV228 | TTTATTATTTATTAAATAATTTATTTATTTAACTAGTAAAGATC CAAATTAGAGG | Sac1_ΔTMD (re) |

**siRNA**

| **NCBI gene** | **Gene description** | **Entrez Gene ID** | **Product name** | **Product ID** |
|---|---|---|---|---|
| Unspecific_AllStars_1 | AllStars | None | Unspecific_AllStars_1 | SI03650318 |
| ARF1 | ADP-ribosylation factor 1 | 375 | Hs_ARF1_10 | SI02757272 |
| SACM1L | SAC1 suppres-sor of actin mutations 1-like | 22908 | Hs_SACM1L_5 | SI02664620 |
| | | | Hs_SACM1L_6 | SI02664627 |
| | | | Hs_SACM1L_8 | SI04438994 |
| | | | Hs_SACM1L_9 | SI04439001 |

[a] Restriction sites and regions overlapping with destination vector are underlined.
